# Cold acclimation and diapause independently improve egg cold tolerance and act through distinct mechanisms in the invasive mosquito *Aedes albopictus*

**DOI:** 10.64898/2026.11.11.765382

**Authors:** Samantha L. Sturiale, Vladimír Košťál, Martin Moos, Nicholas M. Teets, Peter A. Armbruster

**Author notes:** **Corresponding author:** SLS.

## Abstract

1. Phenotypic plasticity is an important mechanism by which organisms can cope with environmental change. However, there is growing recognition that phenotypic plasticity is not a single phenomenon but instead encompasses a range of responses that vary in reversibility and are predicted to evolve under different conditions. Nevertheless, few studies have disentangled the fitness effects and mechanisms of different forms of plasticity, making it difficult to predict their respective roles in promoting population persistence in the face of climate change.
2. In our study, we assessed plasticity of egg cold tolerance in the vector mosquito *Aedes albopictus,* considering both irreversible plasticity caused by photoperiodically-mediated diapause and reversible plasticity caused by cold acclimation. We also analyzed metabolites and lipids to investigate the extent to which these two forms of plasticity improve cold tolerance through shared mechanisms.
3. We found that diapause and cold acclimation independently enhance cold tolerance of *Ae. albopictus* eggs. While the beneficial effect of diapause on cold tolerance persisted throughout our experiment, cold tolerance gained through acclimation was lost when eggs were returned to warm temperatures.
4. Cold tolerance plasticity in *Ae. albopictus* does not appear to be driven by the accumulation of classical cryoprotectants. Instead, reductions in citric acid cycle intermediates and free fatty acids imply that diapause induces metabolic suppression, which likely helps maintain energy balance during winter and may also improve cold tolerance by protecting against cold-induced disruption of metabolic homeostasis. Cold acclimation generally increased lipid unsaturation, which may function to preserve membrane fluidity at low temperatures. Overall, diapause and cold acclimation imparted largely nonoverlapping effects on egg metabolite and lipid profiles, indicating that these two forms of plasticity act through distinct molecular pathways.
5. In addition to identifying potential mechanisms by which an invasive vector mosquito attains cold tolerance to survive winter, our results provide evidence that irreversible and reversible plasticity are distinct phenomena capable of evolving independently during populations’ responses to novel environmental conditions.

## 1) INTRODUCTION

Determining how organisms maximize fitness in heterogeneous environments is a long-standing goal of evolutionary ecology with important implications for anticipating organismal responses to climate change (Bitter et al., 2021; Via et al., 1995). Adaptive phenotypic plasticity is a key mechanism by which organisms cope with environments that vary predictably (Baythavong, 2011; Pfennig, 2021). Plastic responses that cause irreversible changes in phenotype during development (irreversible, fixed, ‘one-shot’, or developmental plasticity) are predicted to evolve in response to variability occurring across generations or at large spatial scales (Botero et al., 2015; Chevin et al., 2010). In contrast, responses that enable reversible shifts in phenotype (i.e., reversible or labile plasticity, acclimation, or phenotypic flexibility) are expected to evolve in response to shorter-term variability that occurs within an individual’s lifetime (Gabriel et al., 2005; Lande, 2014; Piersma & Drent, 2003). While empirical studies have identified cases where a given trait is responsive to both irreversible and reversible plasticity, the relative fitness effects and underlying mechanisms of these different forms of plasticity are rarely disentangled (Stager et al., 2024). This limitation constitutes a key gap in our fundamental understanding of organismal adaptation to environmental heterogeneity.

Characterization of different forms of plasticity and their mechanisms is also necessary to anticipate the vulnerability of species to climate change. First, climate change is generally increasing environmental variability (Kefford et al., 2022), which may negatively impact species relying on irreversible plasticity if this variability occurs within an individual’s lifetime (Stager et al., 2024). Second, climate change is expected to reduce the predictability of environmental variation by altering covariances between proximate environmental cues and the selective environment (Watts et al., 2022), and by increasing the frequency of extreme events (Hansen et al., 2012; Kodra et al., 2011). In an increasingly unpredictable environment, the fitness costs of expressing a mismatched phenotype are anticipated to be particularly strong for irreversible forms of plasticity that do not permit further adjustments to the phenotype (Hoffmann & Bridle, 2022). Importantly, the ability to modify phenotypes using both irreversible and reversible plasticity simultaneously and/or using multiple cues could reduce the negative fitness impacts of mismatch (Beaman et al., 2016; Watts et al., 2022). Finally, responses to climate change on longer timescales will likely involve evolutionary adaptation in plasticity (Bradshaw & Holzapfel, 2001). However, if irreversible and reversible plasticity modify a trait through the same molecular pathways, the resulting genetic correlation between these responses could constrain their further evolution (Chevin & Lande, 2015; Walsh & Blows, 2009).

In arthropods, diapause and cold acclimation are forms of irreversible and reversible plasticity, respectively, that often impact cold tolerance. Diapause is an anticipatory, pre-programmed developmental arrest typically triggered by photoperiodic cues. In some species, cold tolerance is enhanced during diapause independently of winter temperatures (Denlinger, 2002; Lehmann et al., 2018). Cold acclimation is a direct response to nonlethal low temperatures over the course of hours to weeks that increases survival under colder conditions (Fields et al., 1998; Sørensen et al., 2015). Cold acclimation is often reversible and lost when temperatures warm, which may allow organisms to adjust cold tolerance levels continually (Isobe et al., 2013; Šlachta, Vambera, et al., 2002). However, few studies have quantified this reversibility directly. Importantly, the effects of diapause and cold acclimation on cold tolerance are often studied in isolation or confounded by study design (Denlinger, 2022; Hodkova & Hodek, 2004). As a result, the relative contributions of diapause and cold acclimation to cold tolerance, as well as what mechanisms underlie these different forms of plasticity remain unclear. It is critical to address these knowledge gaps to anticipate organismal responses to winter climate change. Specifically, winter temperatures are expected to become more variable and unpredictable due to a reduction in insulating snow cover and an increase in the frequency of extreme events (e.g., winter heatwaves and cold snaps) (Kodra et al., 2011; Williams et al., 2015). Additionally, increasing fall temperatures and poleward range shifts could interfere with the proper timing of diapause induction, meaning that arthropods could increasingly experience cold outside of their overwintering stages (Bale & Hayward, 2010).

In this study, we investigated the mechanisms underlying irreversible versus reversible plasticity by quantifying the effects of diapause and cold acclimation on cold tolerance in eggs of the invasive disease vector, *Aedes albopictus*. In the temperate areas of its worldwide range, this species survives winter by entering embryonic diapause at the egg stage in response to maternal photoperiod cues (Armbruster, 2016). Cold tolerance of *Ae. albopictus* eggs increases in response to both diapause and cold acclimation (Hanson and Craig, 1994), making it an ideal system for testing the relative fitness effects and mechanisms of these different forms of plasticity. We first compared cold tolerance of eggs exposed to a full-factorial combination of diapause and acclimation treatments to quantify the effects of each form of plasticity on cold tolerance and determine the extent to which cold acclimation is reversible. We then used metabolite and lipid analyses to test two prominent mechanisms underlying arthropod cold tolerance.

First, we expected that diapause and cold acclimation would increase levels of cryoprotectants, which are amino acids, sugars, and sugar alcohols that prevent cold injury by facilitating super-cooling and/or stabilizing proteins and membranes (Teets et al., 2023). Second, we predicted that diapause or cold acclimation would cause membrane alterations such as increases in fatty acid unsaturation, decreases in fatty acid length, and increases in the ratio of phosphoethanolamine relative to phosphatidylcholine phospholipid classes. These alterations may improve membrane function at low temperature by increasing membrane fluidity (Sinensky, 1974) and/or by preventing unregulated phase transitions of the phospholipid bilayer (Hazel, 1997). We also determined the degree to which these distinct forms of plasticity have overlapping effects on metabolite and lipid profiles. Our results demonstrate how physiological studies can be leveraged to address fundamental questions in evolutionary ecology, such as the relationship between irreversible and reversible plasticity and the ability of organisms to adapt to climate change.

## 2) MATERIALS AND METHODS

### 2.1) Population origin and animal rearing

The lab colony of *Ae. albopictus* was established using 302 field-caught larvae and pupae from Manassas, Virginia on July 22^nd^, 2021. Individuals were collected from 10 discarded tires at an auto-salvage yard (38°38’04.4“N 77°25’26.8“W). The colony was propagated for 10 generations (minimum adult census population size of ca. 1000 per generation) according to previously described husbandry procedures in a walk-in incubator held at 21°C, 80% relative humidity (RH), and a long-day (LD) photoperiod (16:8 Light:Dark) (Sturiale and Armbruster 2023). The F_10_ generation used to generate eggs for this study was reared under the same conditions in groups of 200 larvae in separate 5.5-L Sterlite containers with 2-L of deionized (DI) water. Upon pupation, individuals were distributed into adult cages (ca. 500 per cage) and kept at 21°C and 80% RH. Half of these cages were placed in the walk-in incubator at LD photoperiod to stimulate production of non-diapause eggs, while the other half were placed inside photoperiod-cabinets set to short-day (SD) photoperiod (8:16 Light:Dark) located in the walk-in incubator (previously described in Mushegian et al. 2021) to stimulate production of diapause eggs. Adult females were blood-fed to produce eggs, which were collected on strips of unbleached paper towel (i.e., egg papers) as described previously ((Sturiale & Armbruster, 2023), supplementary information section 1).

To confirm that LD and SD treatments induced production of non-diapause and diapause eggs, respectively, we measured diapause incidence of eggs (as described previously in Batz et al. 2020) in 30 replicate groups (15 LD and 15 SD), each containing an average of ∼36 eggs (range: 20-65) from a total of five LD and four SD cages (supplementary information section 2).

### 2.2) Egg acclimation treatment groups

To test the effect of acclimation temperature and duration on egg cold tolerance and biochemical profiles, we exposed diapause and non-diapause eggs to different acclimation treatments in a full-factorial design. For measurements of cold tolerance, we used five acclimation treatments: before acclimation, warm acclimation 15-d, cold acclimation 15-d, cold acclimation 22-d, and de-acclimation 22-d; details are provided in Table 1 and Figure 1A, B. For comparisons of biochemical profiles, we used three acclimation groups for metabolomic and lipidomic assays: before acclimation, warm acclimation 15-d, cold acclimation 15-d. We waited until eggs were 11-13 days post-oviposition before initiating the acclimation period to ensure that embryogenesis was completed (Reynolds et al., 2012). After embryogenesis, non-diapause individuals remain as quiescent pharate larvae inside the egg chorion until exposed to water as a hatching stimulus. Diapause individuals also remain as pharate larvae inside the egg chorion but are insensitive to hatching stimulus until diapause has terminated, which takes approximately 100-150 days at 21°C (Batz et al., 2020). Thus, while individuals used for this study are most precisely identified as “pharate larvae within eggs”, we use “eggs” hereafter. All exposures to 21°C took place in the walk-in incubator, while exposures to 10°C and 3.5°C took place in separate temperature- and light-controlled growth chamber (model I-30 VL, Percival Scientific, Perry, IA, USA). Acclimation treatments for non-diapause eggs were all performed at LD photoperiod, while acclimation treatments for diapause eggs were all performed at SD photoperiod.

**Figure 1.**
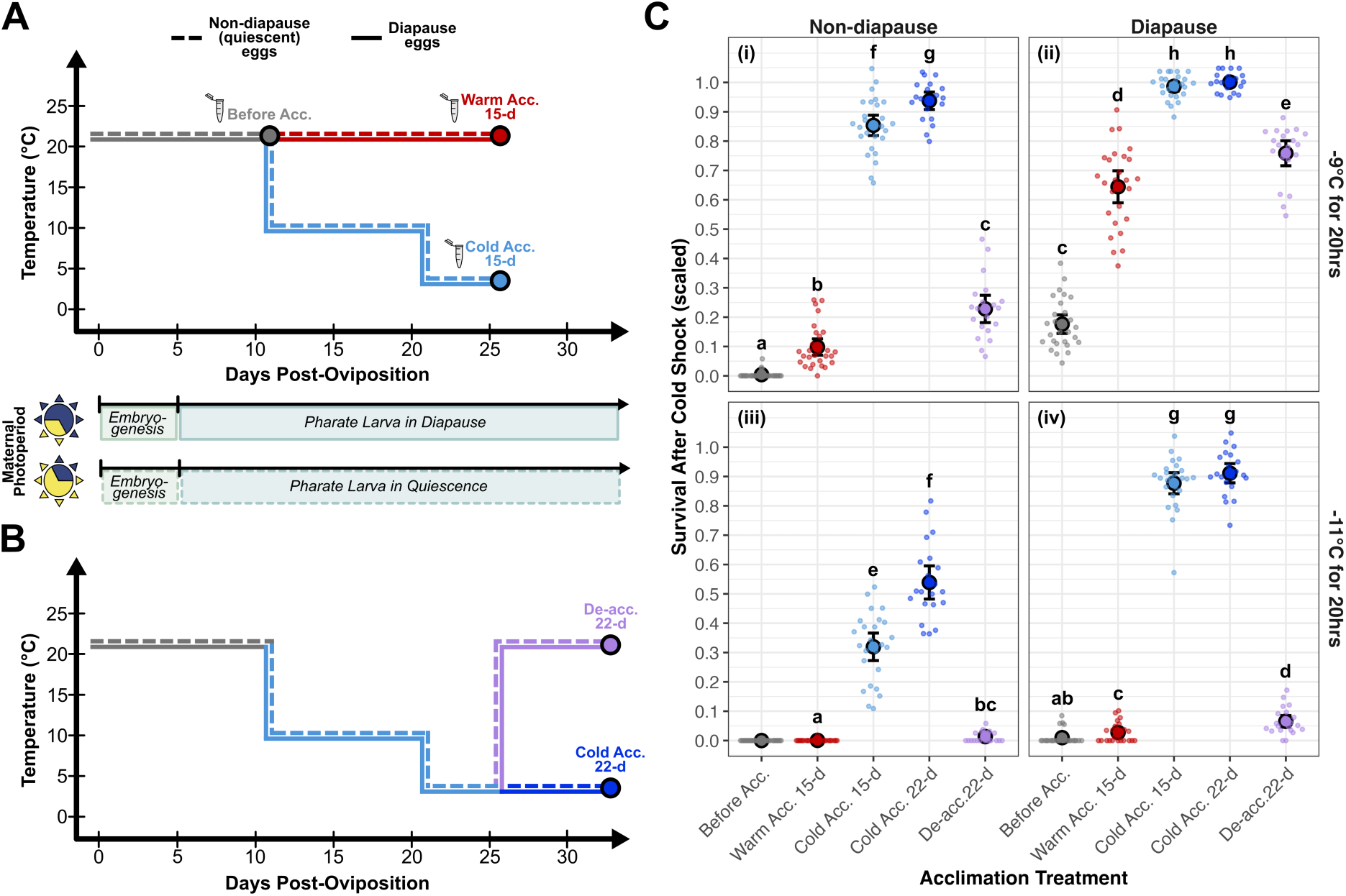
Diapause and cold acclimation independently enhance cold shock survival of *Ae. Albopictus* eggs. Age and temperature exposures for the treatment groups used to test the effects of cold acclimation **(A)** and the reversibility of cold acclimation **(B)**. Egg developmental timeline in the center shows the transition from embryogenesis to pharate larva within diapause or quiescent eggs depending on maternal photoperiod. Circles indicate treatments that were tested for cold shock survival, microcentrifuge tubes indicate treatments that were sampled for metabolite and lipid analysis. **(C)** Survival of each treatment group after exposure to a cold shock of −9°C (i, ii) or −11°C (iii, iv) for 20 hours. Large points and error bars indicate the mean survival ± two standard errors, smaller points show the proportion of surviving individuals in each replicate. Cold shock survival of each replicate was scaled according to the mean hatch rate of the control (non-cold shocked) samples exposed to the same diapause and acclimation treatment. Points with different letters are significantly different (*p*-adjusted < 0.05) based on post-hoc testing (compared separately for −9°C and −11°C cold shocks). Points without letters could not be used in contrasts because all replicates for that treatment group had zero values, leading to complete separation.

**Table 1.**
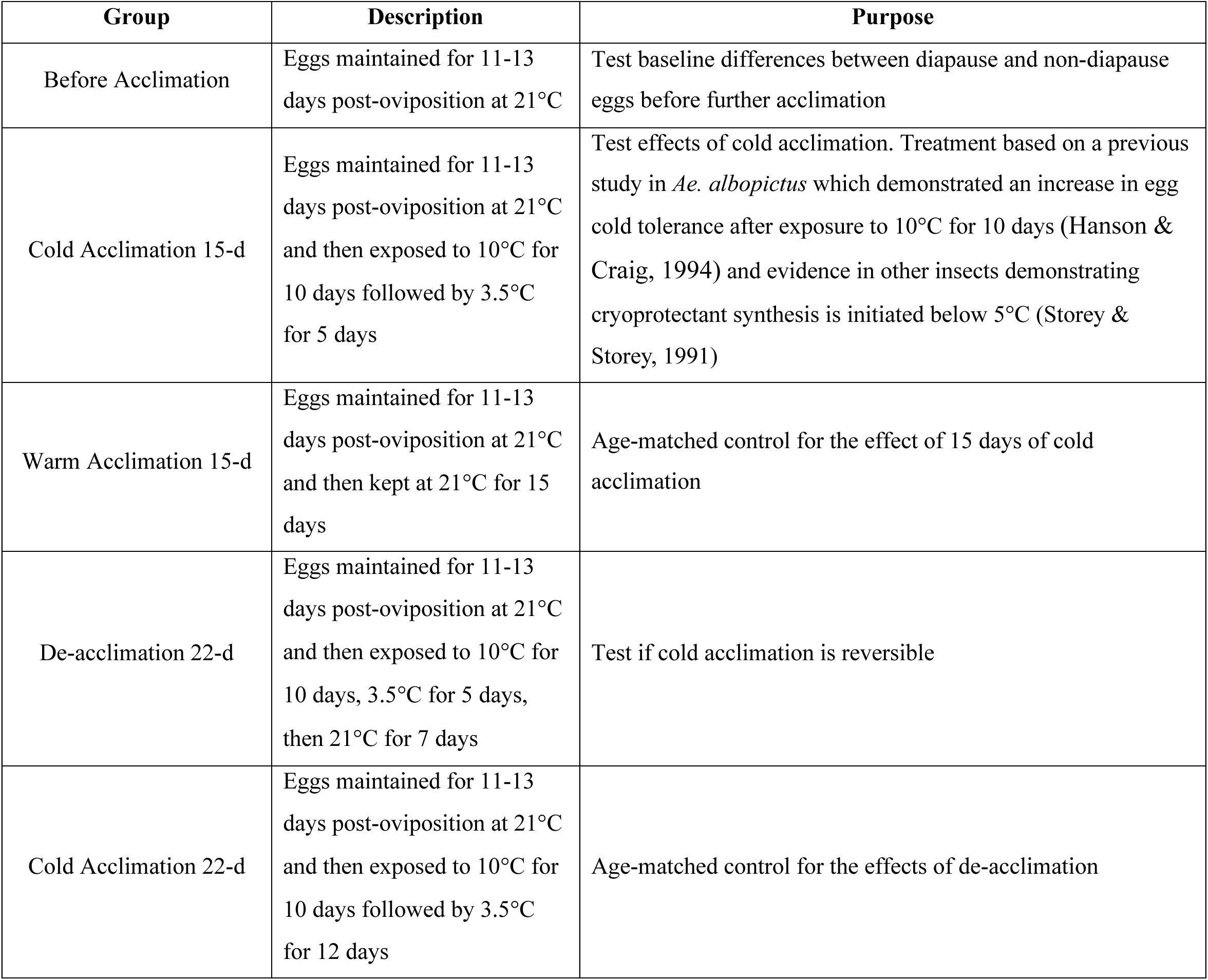
Descriptions of the five acclimation regimes. We exposed both diapause and non-diapause eggs to each acclimation treatment.

### 2.3) Measurement of egg cold tolerance

To test the effects of diapause and acclimation treatments on egg cold tolerance, we conducted acute cold shock trials on eggs from each of the ten diapause and acclimation treatment combinations (Figure 1A, B). Prior to initiation of the acclimation treatments, eggs used in cold shock trials were separated into smaller replicate papers containing ∼36 (range: 20-81) viable (i.e., non-desiccated) eggs. Each replicate paper was placed individually in a 60-mm plastic Petri dish on two layers of filter paper (Fisher Scientific Fisherbrand™, No. 09-795H) which was moistened with 100-μL of DI water to prevent desiccation.

These replicate papers were then assigned to one of five acclimation treatment groups (Table 1, Figure 1A, B). For all acclimation treatments, 60-mm Petri dishes containing individual replicate papers were stored in 2.5-g plastic storage containers (Rubbermaid™) containing a mesh-covered cup with 75-mL of DI water to maintain ∼95% RH.

At the end of acclimation treatments, replicate papers were subjected to one of three possible cold shock treatments to assess cold tolerance: control (no cold shock), −9°C cold shock for 20 hours, and - 11°C cold shock for 20 hours. These cold shock temperatures and durations were chosen because previous studies indicate that the lower lethal temperature for 24 hours of exposure falls between −8°C and −12°C for *Ae. albopictus* eggs of different diapause and acclimation treatment combinations (Hanson & Craig, 1994). Cold shock trials were performed using a programmable refrigerated bath (Thermo Scientific Neslab RTE7) containing a 1:1 mixture of water and ethylene glycol. Following cold shock trials, eggs were stimulated to hatch six times to measure survival of each replicate paper (survival = total number of live hatched larvae divided by the number of eggs on the replicate paper). For a full description of the cold shock trial and hatching protocols, see supplementary information section 3.

A total of seven LD and seven SD parental cages generated eggs that were used for the cold shock trials. The sample size for cold shock trials was 727 replicate egg papers, with an average of 24 (range: 20-31) replicate papers per each combination of diapause treatment, acclimation treatment, and cold shock treatment (see Table S1 for replication statement and Tables S2, S3 for further details).

### 2.4) Preparation of egg samples for metabolite and lipid analyses

To investigate the mechanisms by which *Ae. albopictus* eggs attain cold tolerance following diapause and cold acclimation, we generated diapause and non-diapause eggs and exposed them in a full-factorial manner to three acclimation treatments (before acclimation, warm acclimation 15-d, and cold acclimation 15-d, Figure 1A), producing six total treatment groups. Two sets of 36 samples (six biological replicates for each of the six treatments) were collected, one set for metabolite analysis and the other for lipid analyses. Each sample contained an average of 6.13 mg (range: 2.08-12.49) eggs. Egg samples used for metabolite and lipid analyses were derived from six LD adult cages and six SD adult cages, with each cage contributing one egg sample to each acclimation treatment for both metabolite and lipid sample sets (Table S1). See supplementary information section 4.1 for details of sample preparation, homogenization, and extraction.

### 2.5) Metabolite analysis

We performed relative quantification of 50 select metabolites using a liquid chromatography – mass spectrometry (LC-MS) platform as described previously (Moos et al. 2025, supplementary information section 4.2). These 50 metabolites were chosen for relative quantification due to their roles in metabolic pathways previously implicated in responses to diapause and cold acclimation in other insects (Moos et al. 2025). See Table S4 for the list of metabolites, including abbreviations. If identified metabolites passed quality control checks and showed no contamination of blank samples, relative quantities were determined across samples by measuring the area under chromatographic peaks. The relative peak area of each metabolite was normalized by the sample fresh mass (mg). In downstream analyses, we only included the 45 metabolites which were detected in all samples, but the full dataset is available in supplementary file 1.

### 2.6) Lipid analyses

Extracted lipid samples were used to perform relative quantification of phospholipid (PL) classes as well as absolute quantification (i.e., molar proportions) of fatty acid methyl esters (FAMEs) across the whole sample (rather than identifying FAMEs present in each PL class) (see supplementary information sections 4.3, 4.4, 4.5 for details). To separate and identify PL classes, we used liquid chromatography coupled to high-resolution mass spectrometry, with slight modifications of methods used previously (Cífková et al., 2012). For each sample, peak areas of all lipids in each PL class were summed, and relative abundance was expressed as a percentage of the total. The composition of FAMEs was analyzed by gas chromatography coupled with mass spectrometry, following a slight modification of a previously used method (Vodrážka et al., 2025). FAMEs were expressed as molar percentages of total FAME content.

Fatty acid names are given as X:YnZ, where X is the number of carbons, Y is the number of double bonds, and Z indicates the position of the first double bond from the methyl end of the molecule, if known.

From data on individual PL classes and FAMEs, we calculated four summary variables previously described in studies of arthropod phospholipid composition (Slotsbo et al., 2016): the ratio of phosphoethanolamine to phosphatidylcholine (PE:PC ratio); the unsaturation index (Σ (1×% monoenes + 2×% dienes + 3×% trienes + 4×% tetraenes + 5×% pentaenes)/100); the proportion of poly-unsaturated FAME (% dienes + % trienes + % tetraenes + % pentaenes); and the average FAME length.

### 2.7) Statistical analyses

All statistical analyses and data visualization were performed in R (v4.5.2), and scripts are available on Github (https://github.com/samstur/Albo_Cold_Mechanisms/blob/main/README.md). Additional details are available in supplementary information section 5. For all models described below, we used a type II Anova (car R package, v3.1.5; (Fox and Weisberg, 2019)) to test for significance of fixed effects. Following the detection of a significant fixed effect, we performed post-hoc pairwise comparisons using the emmeans function (emmeans R package v2.0.2; (Lenth and Piaskowski, 2026)), with *p*-values adjusted to account for multiple comparisons (Benjamini and Hochberg 1995).

We tested the effects of diapause and acclimation treatments on cold shock survival using generalized linear mixed effects models (glmmTMB function of glmmTMB R package, v1.1.14; (Brooks et al., 2017)) with a beta-binomial distribution to account for overdispersion and a logit link (Demétrio et al., 2014). A separate model was fit for the eggs exposed to a cold shock temperature of −9°C versus those exposed to −11°C. For both models, diapause treatment, acclimation treatment, and the interaction term were included as fixed effects. Additionally, in both models, egg age at the start of acclimation treatments was included as a covariate and parental cage was included as a random effect.

To visualize overall differences in metabolite, PL class, and FAME profiles, we performed three separate principal component analyses (PCAs). For all three datasets, we performed the PCA with the prcomp function (R stats package, v4.5.2) on data that was scaled and mean-centered using the scale function (R base package, v3.6.2). To test whether treatment groups differed significantly in PC1 and PC2 scores from each PCA, we performed separate Kruskal-Wallis tests (stats R package) followed by pairwise Wilcoxon rank sum tests, with *p*-values adjusted for multiple testing across pairwise comparisons (Benjamini & Hochberg, 1995). Additionally, to test whether variation in overall metabolite, PL class, or FAME profiles was associated with variation in cold tolerance across treatment groups, we tested for correlations between the first two PCs and the average scaled −9°C cold shock survival of each treatment group using either a Pearson or Kendall rank correlation test (cor.test in R stats package). Survival following a −11°C cold shock was not used in correlation tests because multiple treatment groups had zero or near-zero values for this cold shock treatment.

To test for significant effects of diapause and cold acclimation on individual metabolites, PL classes, and FAME species, we ran a separate linear mixed effect model (lmer function in lme4 R package, v2.0.1; (Bates et al., 2015)) for each of the response variables across these three datasets. In each model, total treatment (i.e., the combination of diapause and acclimation treatments) was included as a fixed effect and parental cage was included as a random effect. Following the detection of a significant effect of total treatment, we performed a planned set of five post-hoc pairwise contrasts that were of biological interest to test for effects of cold acclimation (Figure 3A, contrasts 1-2) and the effects of diapause (Figure 3A, contrasts 3-5). In some cases, data failed to meet the assumption of normality of residuals, so we instead used non-parametric tests (see supplementary information section 5). For each of the five pairwise contrasts we also computed log_2_ fold-change (log_2_FC). In contrasts which describe an effect of cold acclimation, a positive log_2_FC indicates higher values in cold-acclimated samples relative to warm-acclimated samples. In contrasts which describe an effect of diapause, a positive log_2_FC indicates higher values in diapause samples relative to non-diapause samples.

Finally, to test for differences in lipid summary variables across the six total treatments, we used separate Kruskal-Wallis tests followed by pairwise Wilcoxon rank sum tests, with *p*-values adjusted for multiple testing across pairwise comparisons (Benjamini & Hochberg, 1995).

## 3) RESULTS

### 3.1) Cold shock survival

Survival of control (non-cold-shocked) eggs was high across treatments (mean: 0.95, range: 0.92-0.97, Figure S1). Cold shock survival after exposure to −9°C was significantly affected by diapause treatment (χ^2^_1, 234_ = 319.21, *p* < 0.001), acclimation treatment (χ^2^_4, 234_ = 1247.91, *p* < 0.001), and the interaction (χ^2^_4, 234_ = 16.14, *p* = 0.003) (Figure 1C, panels i-ii). Based on post-hoc testing (Table S5), both diapause and non-diapause eggs had significantly greater survival after undergoing cold acclimation for 15 days, compared to eggs tested before acclimation and age-matched warm acclimated eggs (all *p* < 0.001). Non-diapause eggs exposed to a longer cold acclimation period (22 days) had greater survival compared to eggs cold-acclimated for 15 days (*p* < 0.001), with no significant difference between these two acclimation groups in diapause eggs (*p* = 0.213). In both non-diapause and diapause groups, eggs exposed to 15 days of cold acclimation followed by seven days at warm temperature (de-acclimation group) had significantly lower survival than the age-matched group exposed to cold acclimation for 22 days (both *p* < 0.001). Finally, we observed that eggs exposed to warm acclimation for 15 days exhibited greater survival than eggs in the before acclimation group, particularly for diapause eggs (both *p* < 0.001). Across all acclimation treatments, diapause eggs exhibited greater cold shock survival than non-diapause eggs (all *p* < 0.001).

Survival following cold shock at −11°C was lower than survival after −9°C exposure, and it was significantly affected by diapause treatment (χ^2^_1, 173_ = 224.89, *p* < 0.001) and acclimation treatment (χ^2^_3, 173_ = 1111.14, *p* < 0.001), but not their interaction (χ^2^_3, 173_ = 6.87, *p* = 0.076) (Figure 1C, panels iii, iv). Patterns of −11°C cold shock survival across groups were largely similar to those in the −9°C cold shock survival data (see Table S6 for results of post-hoc tests).

### 3.2) Metabolite profile

In the PCA of diapause and cold acclimation effects on metabolite profiles, samples largely separated according to diapause treatment on PC1 (35.9% of variation) and acclimation treatment on PC2 (18.2% of variation) (Figure 2A, Figure S2). PC1 was not correlated with survival following cold shock at −9°C (r = 0.032, *p* = 0.852, Figure 2B), while PC2 exhibited a positive correlation with −9°C survival (τ = 0.542, *p* < 0.001, Figure 2C). PC1 loadings show that diapause samples (negative PC1 values) were characterized primarily by lower levels of glycolysis and citric acid cycle intermediates (e.g., glucose-6-phosphate, lactic acid, pyruvic acid, etc.) (Figure 2D). PC2 loadings indicate that cold acclimation samples (positive PC2 values) were characterized by lower levels of most metabolites (Figure 2E).

**Figure 2.**
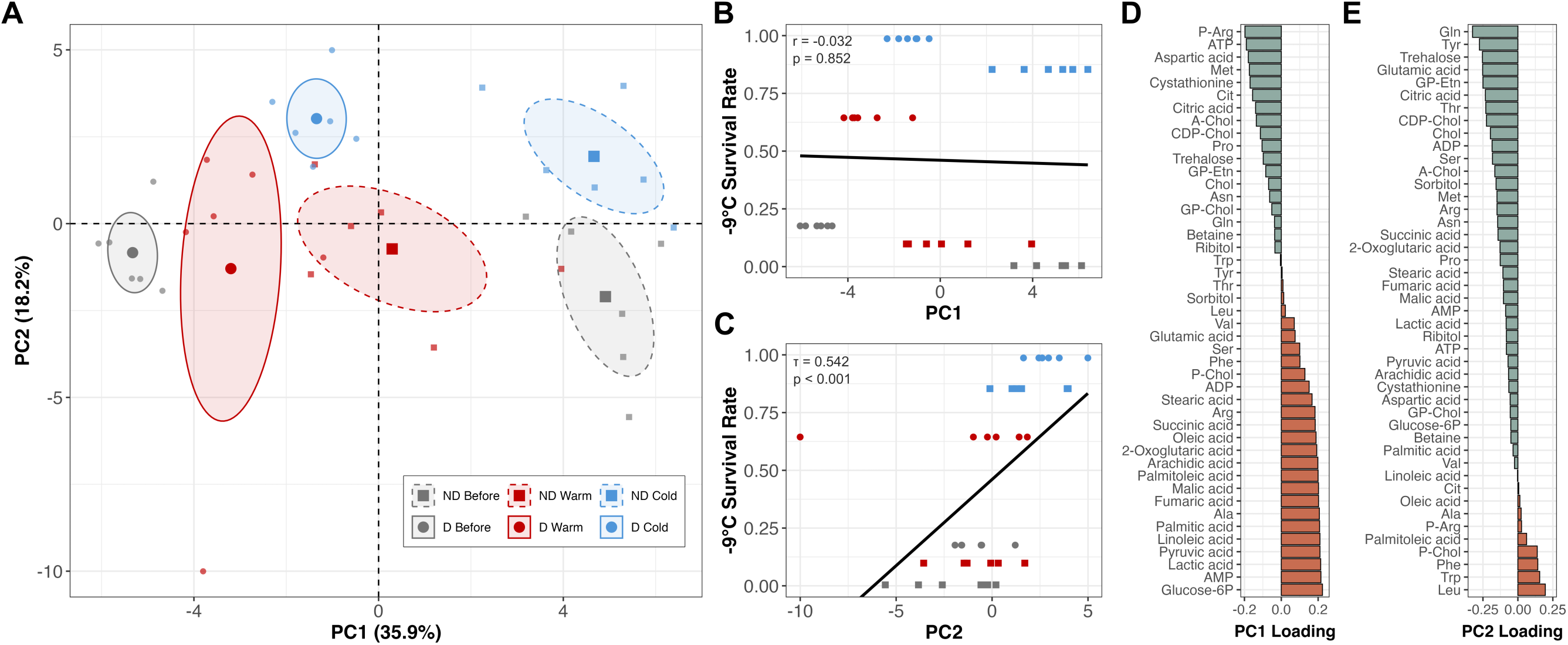
Both diapause and acclimation affect overall metabolite profiles. **(A)** Principal component analysis (PCA) plot showing separation of egg samples based on variation in metabolite profiles. Each individual sample is shown as a small point, with the group mean and 95% confidence interval represented by a larger point surrounded by an ellipse. Correlations (Pearson coefficient in **B** and Kendall rank coefficient in **C**) between the first two PC dimensions and the average −9°C cold shock survival of each treatment group are shown in **(B)** and **(C)**, with color and shape formatting as indicated by the legend in **(A)**. Metabolites contributing to variation in PC1 **(D)** and PC2 **(E)** ranked according to their loading value (positive values as orange and negative values as green). Treatment abbreviations are as follows: ND = non-diapause; D = diapause; Before = before acclimation; Warm = warm acclimation 15-d; Cold = cold acclimation 15-d).

Examination of individual metabolites using a planned set of five pairwise contrasts indicated that few metabolites changed in the same direction in response to cold acclimation and diapause (Figure 3A, B; Supplementary File 1). Several metabolites were lower in response to cold acclimation in both non-diapause and diapause eggs (bolded labels in Figure 3C), including polyols (ribitol, sorbitol), amino acids (tyrosine, betaine), and metabolites involved in glycerophospholipid metabolism (choline, acetylcholine, CDP-choline). Only phospho-choline and alanine increased under cold acclimation conditions in both diapause and non-diapause eggs. Diapause eggs exhibited higher phenylalanine, valine, and leucine after cold acclimation (Figure 3C, right panel). The metabolites with lower levels in diapause relative to non-diapause eggs across all three acclimation groups (Figure 3D, bolded labels) included several glycolysis and citric acid cycle intermediates (lactic acid, pyruvic acid, fumaric acid, succinic acid) and free fatty acids (arachidic acid, linoleic acid, stearic acid).

**Figure 3.**
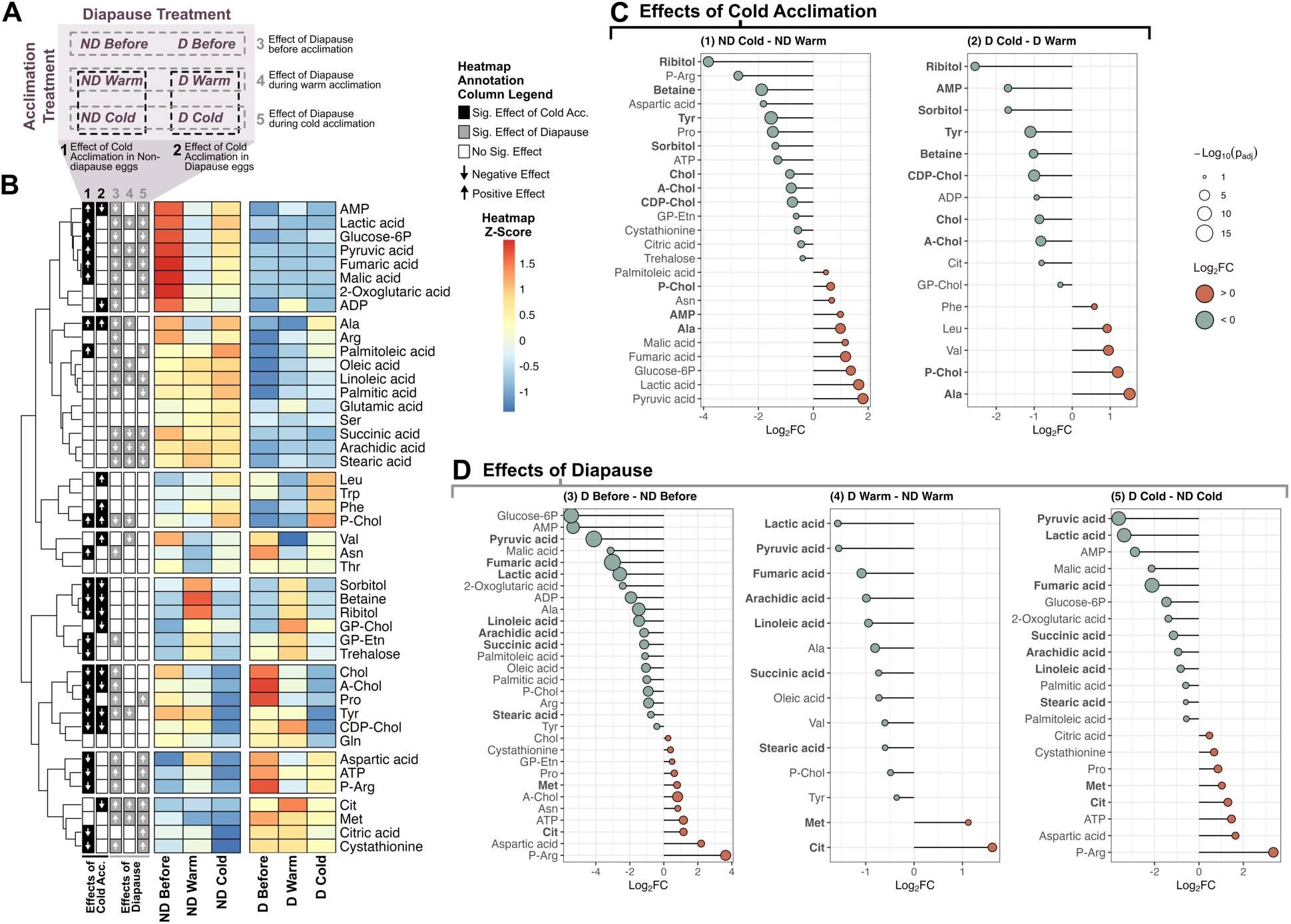
Distinct metabolite shifts due to cold acclimation and diapause. **(A)** Schematic showing the five pairwise contrasts used to test for the effects of cold acclimation (contrasts 1-2) versus the effects of diapause (contrasts 3-5) across the six total treatment groups. **(B)** Heatmap of metabolites clustered according to their similarity in relative abundance (z-score of mass-scaled peak area) across the six treatment groups (averaged across the six replicates within each group). Annotation columns on the left of the heatmap show whether each metabolite was significantly different (adjusted *p* <0.05) across each of the five pairwise contrasts described in **(A)**. Lollipop plots illustrate the log_2_ fold-change (log_2_FC) and - log_10_(*p*-adj.) values of metabolites which differed significantly across each of the five pairwise contrasts, with **(C)** showing pairwise contrasts associated with cold acclimation (contrasts 1-2) and **(D)** showing pairwise contrasts associated with diapause (contrasts 3-5). Metabolites in bold are those which differ significantly across either both pairwise contrasts related to cold acclimation **(C)** or across all three pairwise contrasts related to diapause **(D)**. Treatment group abbreviations are as in Figure 2.

### 3.3) Phospholipid classes and fatty acid methyl ester compositions

We observed limited separation among treatment groups in PCAs of phospholipid (PL) classes and fatty acid methyl ester (FAME) species (Figures S3A, S4A). However, we detected significant positive correlations between average −9°C cold shock survival and PC2 for PL class (*r* = 0.625, *p* < 0.001, Figure S3E) as well as PC1 for FAME PC1 (*r* = 0.53, *p* < 0.001, Figure S4D). We assessed whether cold acclimation and diapause impacted individual PL classes or FAME species using the same five pairwise contrasts as in the metabolite data (Figure 4A, B; Supplementary File 1). Several PL classes were significantly affected by cold acclimation, but largely only in non-diapause eggs (contrast 1, Figure 4A). Diapause also affected most PL classes, but largely only in eggs in the before and/or warm acclimation treatments (contrasts 3 and 4) (Figure 4A). For FAME species, only non-diapause eggs showed effects of cold acclimation (contrast 1) and only one FAME species (C18.3n6) was impacted by diapause (Figure 4B). See Tables S7, S8 for mean ±2 SE values for PL classes and FAME species in each treatment group.

**Figure 4.**
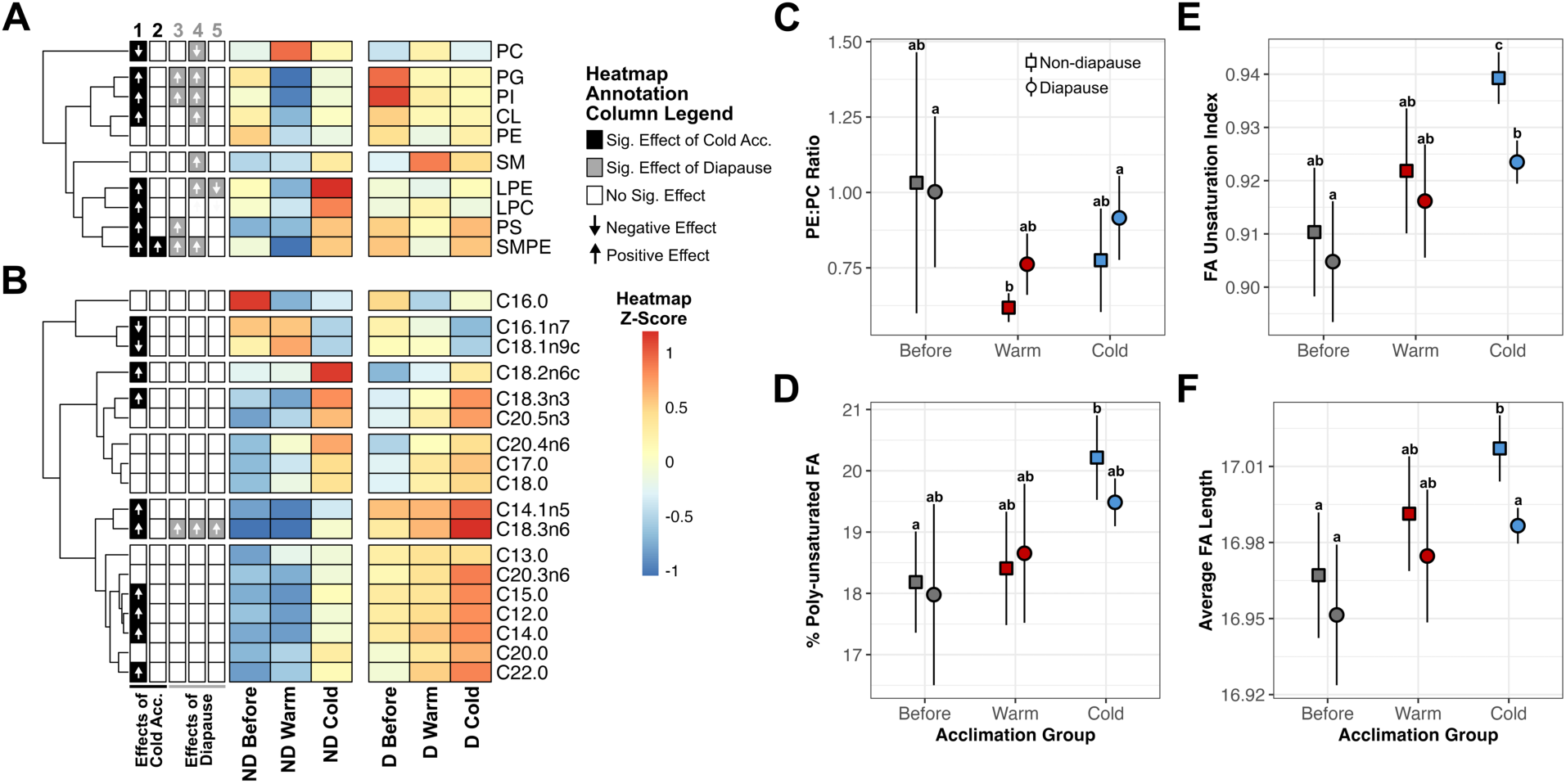
Lipid membrane composition shows an overall stronger signature of cold acclimation than diapause. Heatmaps of phospholipid (PL) classes **(A)** and of fatty acid methyl ester (FAME) species **(B)** clustered according to their similarity in relative abundance across the six total sample groups. Heatmap annotation columns, symbols, conventions, and treatment group abbreviations as in Figure 2. Panels **(C)-(F)** show the mean ± two standard errors of each treatment group for four lipid summary variables: PE:PC phospholipid class ratio, fatty acid (FA) unsaturation index, total proportion of poly-unsaturated FA, and average FA length. Points with different letters are significantly different (*p*-adjusted < 0.05) in post-hoc testing.

All lipid summary variables differed significantly across treatment groups (Table S9). Based on post-hoc testing (Table S10), cold acclimation did not significantly increase PE:PC relative to the before-or warm acclimation treatment groups in either diapause or non-diapause eggs (Figure 4C). In non-diapause eggs, the FA unsaturation index was higher in the cold acclimated group compared to both before acclimation (*p* = 0.011) and warm acclimation groups (*p* = 0.038). In diapause eggs, FA unsaturation index of the cold acclimation group was only significantly greater than that of the before acclimation group (*p* = 0.013) (Figure 4D). Neither the percentage of poly-unsaturated FAs nor the average FA length differed significantly across acclimation groups in diapause eggs (all *p* > 0.05) (Figure 4E, F). In non-diapause eggs, the cold acclimation group exhibited a greater percentage of poly-unsaturated FAs (Figure 4E) and a longer average FA length (Figure 4F) than the before acclimation group (*p* = 0.032 and *p* = 0.043, respectively) but not the warm acclimation group (both *p* < 0.05). Both the FA unsaturation index and average FA length of non-diapause cold-acclimated eggs was higher than that of diapause cold-acclimated eggs (*p* = 0.013 and *p* = 0.043, respectively), but otherwise there were no significant differences in lipid summary variables based on diapause.

### 3.4) Comparison of cold acclimation and diapause responses

Overall, relatively few features (i.e., metabolites, PL classes, and FAME species) showed consistent responses to diapause and cold acclimation (Figure 5, red bars). For example, of the features that significantly increased in response to diapause and/or cold acclimation (i.e., exhibited a positive log_2_FC; Figure 5A), 20 were increased exclusively by cold acclimation (black bars) and 12 were increased exclusively by diapause (gray bars). In contrast, only eight features exhibited positive effects of both diapause and cold acclimation across all incomplete combinations of the five pairwise contrasts examined (Figure 5A, red bars). Similarly, in features showing a significant negative effect (i.e., negative log_2_FC; Figure 5B), only four exhibited decreases in response to both diapause and cold acclimation (red bars).

**Figure 5.**
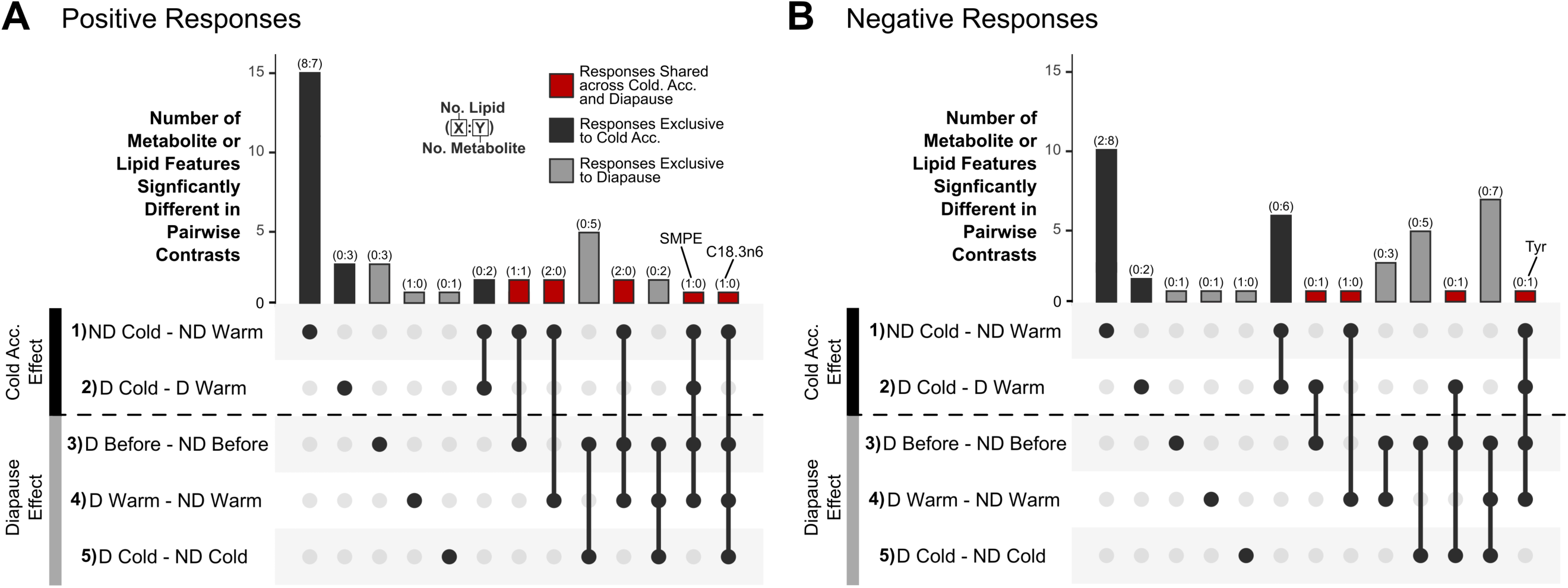
Limited overlap in the metabolite and lipid features that change in response to cold acclimation and diapause. Panel **(A)** shows overlap in features with a positive log_2_FC (i.e., higher in response to either cold acclimation or diapause) while panel **(B)** shows overlap in features with a negative log_2_FC (i.e., lower in response to either cold acclimation or diapause) via UpSet plots. For each panel, the top histogram plot illustrates the number of features that differ significantly in one or more of the five pairwise contrasts. Dots below each histogram bar indicate which pairwise contrasts sets are included in the intersection and thus overlap in significantly different features. The dotted black line in the lower portion of each panel indicates the boundary between the effects of cold acclimation and the effects of diapause. Any intersection that crosses this boundary describes shared significant responses to cold acclimation and diapause (red bar), while any intersection that does not cross this boundary describes responses exclusive to either cold acclimation (black bar) or diapause (gray bar). Text labels in the top plot of each panel identify features that show consistent effects across at least four pairwise contrasts. The numbers above each histogram bar indicate the number of features represented in each bar. Sample group abbreviations correspond with those in Figure 2.

## 4) DISCUSSION

To determine how overwintering arthropods may be impacted by increasing environmental variability and unpredictability due to climate change, it is important to quantify the degree to which cold tolerance is influenced irreversibly through photoperiodic diapause and reversibly through cold acclimation. In our results, we found that diapause and cold acclimation both independently enhanced *Ae. albopictus* egg cold tolerance and exerted cumulative benefits when combined (Figure 1C), consistent with a past study in this species (Hanson & Craig, 1994). We further show for the first time in *Ae. albopictus* that cold acclimation was reversible in response to warming while diapause-associated cold tolerance was retained over the course of the experiment (Figure 1C). Few studies have investigated diapause and cold acclimation in a full-factorial manner to identify their individual and combined effects on cold tolerance (Hodkova & Hodek, 2004). Similar to our results, previous work largely indicates dual regulation of cold tolerance by both forms of plasticity (but see (Štětina et al., 2025)), but species appear to differ in which plastic response exerts a stronger effect and whether, when combined, the effects of diapause and cold acclimation are redundant or cumulative (Khodayari et al., 2012; Koštál et al., 2001, 2003; Šlachta, Berková, et al., 2002; Šlachta, Vambera, et al., 2002). In our study, cold acclimation improved cold tolerance to a greater degree than diapause (Figure 1C). This result contradicts the assumption that irreversible plasticity should allow for greater phenotypic change compared to reversible plasticity (Hoffmann & Bridle, 2022), potentially altering the expected relative fitness costs of each form of plasticity as environmental unpredictability increases. Additionally, the co-expression of both forms of plasticity is not often considered in modeling studies of the evolution of plasticity but may be beneficial in protecting against phenotype-environment mismatch arising due to climate change (Beaman et al., 2016; Watts et al., 2022). For example, a warming climate may contribute to phenotype-environment mismatch by preventing adequate cold acclimation in early winter or by causing untimely de-acclimation during mid-winter heatwaves (Sobek-Swant et al., 2012). However, diapause-induced cold tolerance may mitigate the disruption of cold acclimation, as supported by the present study (Figure 1C) and the observation that winter heatwaves did not reduce the overwintering survival of diapause *Ae. albopictus* eggs (Sturiale & Armbruster, 2023).

We found limited support for the hypothesis that cold tolerance in *Ae. albopictus* eggs is improved via the accumulation of classical cryoprotectants. For example, the levels of several sugars and sugar alcohols (sorbitol, ribitol, trehalose), which often accumulate seasonally in arthropods (Koštál et al., 2001; Storey & Storey, 2012), were unaffected by diapause and were higher in warm-acclimated eggs (Figure 3b). Further, proline, an amino acid with well-established cryoprotective properties (Grgac et al., 2022; Koštál et al., 2011), was reduced by cold acclimation and exhibited only slight increases in response to diapause (Figure 3b). Cold tolerance plasticity in other arthropods is not always accompanied by large increases in classical cryoprotectants. For example, among 15 cold-hardy *Drosophila* species that possess strong cold tolerance plasticity, most exhibited only moderate or weak cryoprotectant accumulation during acclimation (Moos et al., 2025). Similarly, plasticity on shorter timescales via rapid cold hardening can significantly improve cold tolerance levels but does not always stimulate increases in classical cryoprotectants (Teets et al., 2012; Teets & Denlinger, 2013). In combination with these studies, our results suggest that mechanisms other than cryoprotectant accumulation likely underlie the strong effects of cold acclimation and diapause on egg cold tolerance in *Ae. albopictus*.

Diapause and cold acclimation also induced shifts in several amino acids that have been associated with cold tolerance but are not considered classical cryoprotectants. For example, we observed that phenylalanine (Phe) and leucine (Leu) increased in response to cold acclimation, but only in diapause eggs (Figure 3C). Greater Phe and/or Leu levels have been noted in response to diapause (Michaud & Denlinger, 2007) and cold acclimation (Fields et al., 1998; Lalouette et al., 2007) in other arthropods, and studies in plants demonstrate that these amino acids can mitigate cold-induced damage (Cui et al., 2026; Luo et al., 2025). Additionally, in our study, both diapause and cold acclimation caused reductions in tyrosine (Tyr) (Figures 3C, D). Other arthropods similarly exhibit low Tyr as a result of cold acclimation (Fields et al., 1998; Lalouette et al., 2007; Štětina et al., 2025), diapause (Michaud & Denlinger, 2007), and selection on cold tolerance (Williams et al., 2014). Tyr is a precursor in the synthesis of several insect neurohormones (octopamine, dopamine, tyramine), as well as melanin, a key component of cuticles (Arakane et al., 2016; Roeder, 1999). Thus, decreases in Tyr could reflect changes in neurohormones that serve a protective role during cold shock (Srithiphaphirom et al., 2019) or shifts in egg cuticle ultrastructure that may reduce vulnerability to inoculative freezing (Kreß et al., 2016). Future studies should aim to identify the exact cause of cold shock-induced mortality in this species, as this would clarify the degree to which diapause and cold acclimation modify cold tolerance through mechanisms that mitigate pre-freeze injury versus prevent inoculative freezing.

Another well-described mechanism of cold tolerance plasticity is membrane restructuring that preserves membrane fluidity and/or prevents the transition to a non-functional gel phase membrane at low temperature (Enriquez & Teets, 2025; Kostal, 2010). Consistent with the expectations of this mechanism (Sinensky 1974, Hazel 1997) and studies in other arthropods (Enriquez & Colinet, 2019; Michaud & Denlinger, 2006), we observed that cold acclimation generally increased the FA unsaturation index (Figure 4E). Further, both cold acclimation and diapause influenced several individual PL classes, and cold acclimation alone influenced multiple FAME species (Figure 4A, B). These shifts could be functionally important for cold tolerance given the significant correlations between PC2 for PL class and PC1 for FAME with cold shock survival (Figure S3E, S4D). It is important to note that the position of FAs in the glycerophospholipid molecule can affect their relative influence on overall membrane fluidity (Hazel, 1997). Thus, further examination of PL molecular composition would help to discern whether the shifts that we observed in individual PL classes, individual FAME species, and overall FA unsaturation could meaningfully alter membrane properties and contribute to egg cold tolerance.

We also found evidence that diapause-induced cold tolerance may be related to metabolic suppression. Diapause eggs had lower levels of glycolysis intermediates, citric acid cycle intermediates, free FAs, and ADP/AMP, as well as higher levels of ATP and phospho-arginine relative to non-diapause eggs (Figure 2D, Figure 3D). These patterns, in combination with past transcriptomic and metabolomic data on diapause versus non-diapause eggs of this species, generally imply lower metabolic activity and greater energy storage during diapause (Batz & Armbruster, 2018; Poelchau et al., 2013). Metabolic suppression is a hallmark of diapause (Hahn & Denlinger, 2007), and it is possible that this response provides dual benefits to overwintering arthropods via both energy conservation and protection from cold stress. Specifically, metabolic suppression may reduce the risk of cold-induced disruption of homeostasis between metabolic pathways, thereby preventing possible ATP deficiency, build-up of toxic metabolic byproducts, and loss of ion balance (MacMillan, 2019; Pullin et al., 1991). Consistent with a role for maintenance of metabolic homeostasis in cold tolerance, both selection for cold tolerance and cold acclimation reduce the degree of metabolite responsiveness to cold stress, indicating greater robustness of metabolic networks (Colinet et al., 2012; Enriquez et al., 2018; Williams et al., 2014).

The degree to which populations can adapt to novel climatic conditions through changes in plasticity may be constrained if irreversible and reversible responses act through overlapping mechanisms and thus are genetically correlated. We found that diapause and cold acclimation exerted largely distinct biochemical changes. For example, in the metabolite PCA, diapause and cold acclimation generally drove separation of samples along different axes (Figure 2A). Further, we observed only 12 total instances where a metabolite or lipid species changed in the same direction in response to cold acclimation and diapause, and most of these shifts occurred in only some of the contrasts used to test the effects of each response (i.e., most were not consistent effects of either response) (Figure 5). The observation that diapause and cold acclimation have largely distinct effects on biochemical profiles suggests that these two plastic processes likely act through different molecular pathways, an interpretation that is further supported by the cumulative effects of these responses on cold tolerance when co-expressed (Figure 1C). Of the few previous studies that also examined the biochemical effects of diapause and cold acclimation in a full-factorial manner, one did not directly compare the effects of each response on metabolite profiles (Štětina et al., 2025), some noted overlapping effects on phospholipid composition (Koštál et al., 2003) or on one of five measured cryoprotectants (Koštál & Šimek, 1994), and some observed largely different effects of each response on metabolite profiles (Khodayari et al., 2013; Willot et al., 2026) similarly to our results. Further, a study that separately measured the effects of pupal diapause and adult rapid cold hardening on metabolite profiles concluded that these two responses likely modulated cold tolerance through different mechanisms (Michaud & Denlinger, 2007). Evidence of distinct mechanisms is also found in studies showing different metabolomic effects (Enriquez et al., 2018) or quantitative trait loci (Gerken et al., 2015) associated with long-term versus short-term cold acclimation responses. While more data across species is needed to form generalizations about the degree of mechanistic overlap between plastic responses, our results imply that irreversible and reversible forms of cold tolerance plasticity can evolve independently in *Ae. albopictus*. Such decoupling may allow this species to adapt more efficiently to climate change-driven shifts in environmental variability and predictability that could favor one form of plasticity over another (Stager et al., 2024).

## Conclusions

In our study, we comprehensively manipulated diapause and cold acclimation in *Ae. albopictus* eggs to assess the relative phenotypic effects and possible mechanisms of irreversible and reversible forms of plasticity. We found that both forms of plasticity independently enhance egg cold tolerance, with largely distinct effects on metabolite and lipid profiles. In addition to testing potential cold tolerance mechanisms in this species, our results address a key gap in our fundamental understanding of organismal adaptation to thermal heterogeneity by isolating the roles of different forms of plasticity in the regulation of an ecologically significant trait.

## Notes

https://github.com/samstur/Albo_Cold_Mechanisms

### Competing Interest Statement

The authors have declared no competing interest.

### Funding

- National Institutes of Health: R01A13409
- United States Department of Agriculture National Institute of Food and Agriculture: Hatch Project 700545

