## Supplementary Materials for "Cold acclimation and diapause independently improve egg cold tolerance and act through distinct mechanisms in the invasive mosquito *Aedes albopictus*"

**This file contains:**

Supplementary Information

Figures S1 to S4

Tables S1 to S10

### Supplementary Information

#### 1. Experimental Egg Generation

Between 8-11 days after the last pupae was added, each adult cage was blood-fed to repletion on a human host. The Georgetown University Institutional Review Board (IRB) has determined that mosquito blood feeding is not human research and thus does not require IRB approval; however, the blood feeding protocol has been approved by the Georgetown University Occupational Health and Safety Office. Three to four days after blood-feeding, each adult cage was given a plastic cup lined with a strip of unbleached paper towel (i.e., egg paper) and half-filled with DI water. Beginning five days post-blood-feeding, the egg paper of each adult cage was collected and replaced daily for approximately a week. Egg papers were left moist for three days then gently dried. Dried egg papers were stored in individual plastic 90-mm Petri dishes inside 9.5-L plastic storage containers containing a mesh-covered cup with 75-mL of DI water to maintain approximately 95% RH. Throughout the experiments, non-diapause egg papers were kept in the walk-in incubator at LD photoperiod while diapause egg papers were kept in photoperiod-cabinets inside the walk-in incubator at SD photoperiod, unless otherwise noted.

#### 2. Diapause Incidence Measurement

Diapause incidence was measured as previously described (Batz et al., 2020). Briefly, replicate egg papers were stimulated to hatch in 60-mm Petri dishes with 20-mL of DI water and two drops of dog food slurry at 14 days post-oviposition. Diapause eggs were hatched in SD conditions, while non-diapause eggs were hatched in LD conditions. After three days, the number of hatched larvae were counted, the egg papers were dried, and the hatching protocol was repeated the next day. After the second hatching, the remaining embryos were left in a bleach solution for one day (Trpiš, 1970), and the number of embryonated embryos was noted. Diapause incidence of each replicate was calculated as: (number of embryonated, unhatched eggs)/(number of hatched eggs + number of embryonated, un-hatched eggs). The average diapause incidence of eggs laid by LD-exposed parental cages was 0.040, while average diapause incidence of eggs laid by SD-exposed parental cages was 0.997.

#### 3. Cold Shock Trial and Hatching Protocol

The day before cold shock trials, each replicate paper assigned to either -9°C or -11°C cold shock treatment was placed in a 2-mL clear Eppendorf tube (Fisher Scientific 2.0-mL MCT Graduated Natural Cat No. 05-408-138) within their individual 60-mm Petri dishes. The morning of cold shock trials at ZT0, all tubes were sealed then placed in foam floating tube racks (Fisher Scientific Fisherbrand™, No. 36-

099-2328). For all acclimation groups being kept at 21°C, replicate papers were moved immediately to the freezing bath (Thermo Scientific Neslab RTE7), which was set to 21°C. Starting at ZT0.25, the freezing bath was then manually ramped down 2-2.2°C every thirty minutes. Once the bath reached 3.5°C, replicate papers from all acclimation treatments being held at 3.5°C were added to the bath. Ramping continued at the same rate until the bath reached either -9°C (ZT6.75) or -11°C (ZT7.25). The final temperature was held for 20 hours, then the replicate papers were each removed from their tubes and placed carefully back in their assigned Petri dishes. To externally validate the freezing bath temperature during the cold shock trials, an iButton (DS1925, iButtonLink LLC, Whitewater, WI) was placed in a 30-mL glass vial and submerged to the same depth as the Eppendorf tubes using a retort stand and clamp.

Control (i.e., non-cold shocked) replicate papers of each acclimation treatment were placed directly at either 21° (for non-diapause eggs) or 10°C (for diapause eggs) the day of cold shock trials. Following cold shock trials, cold-shocked non-diapause replicate papers were placed at 21°C and LD photoperiod in the walk-in incubator for one day to allow for recovery, then hatched to determine survival. To stimulate hatching, each replicate paper was put in its Petri dish with 20-mL of DI water and two small drops of dog food slurry as described previously (Sturiale and Armbruster 2023). After three days, the number of living hatched larvae per replicate was recorded and the replicate paper was gently dried and then placed back in its Petri dish lined with moist filter paper. Live larvae were identified as those which moved when touched gently with a pipette tip. Non-diapause eggs were re-hatched again after one day, after three weeks, then approximately monthly for a total of six hatch attempts. For diapause eggs, after the cold-shock, replicate egg papers were placed at 10°C and SD photoperiod in a growth chamber to simulate overwintering for 90 days (as in (Hanson & Craig, 1994, 1995)). This procedure was implemented because the survival of diapause eggs cannot be assessed until diapause is terminated. After overwintering, diapause replicate papers were moved to 21°C and LD photoperiod in the walk-in incubator for two weeks to simulate the arrival of spring conditions. Each replicate paper was then hatched a total of six times following the same schedule as the non-diapause eggs. The final hatching of diapause egg samples was performed when eggs were ca. 230 days post-oviposition, which is substantially beyond the 185 days previously documented for diapause termination in North American *Ae. albopictus* populations (Batz et al., 2020).

### 4. Metabolite and Lipid Profile Measurement

#### *4.1 Sample preparation and homogenization*

As with cold shock replicate papers, eggs used for metabolite and lipid analyses were 11-13 days post-oviposition when sampled (for the before acclimation treatment) or when the 15-day acclimation period was initiated (for warm and cold acclimation treatments). The acclimation treatments for the metabolite and lipid analysis were implemented in the same manner as with eggs used for cold shock trials except that egg papers were not cut into small replicates. Egg papers were thus stored in 90-mm Petri dishes without the need for added moist filter paper. Following the completion of the acclimation treatments (with no exposure to cold shock), each egg paper was checked under the microscope, and any desiccated eggs were removed. Then the remaining viable eggs for each sample were gently brushed off their oviposition paper into a pre-weighed 2-mL microcentrifuge tubes (Eppendorf™ Snap-Cap Microcentrifuge Safe-Lock™Tubes, CAT 05-402-7) using a metal spatula. The tube was weighed to the nearest 0.01 mg on a microbalance (Explorer EX125D, Ohaus, Parsippany, NJ, USA) then snap frozen in liquid nitrogen and stored at -80°C. These samples were all frozen between ZT6.5 and ZT7.5. On the day that samples were shipped to a separate lab for extraction and analysis, 400 microliters of 2:2:1 methanol:acetonitrile:water (Methanol, Optima™ LC/MS Grade, Fisher Chemical, CAT A456-1; Acetonitrile, Optima™ LC/MS Grade, Fisher Chemical™, CAT A955-500; Water, Optima™ LC/MS Grade, Fisher Chemical™, CAT W6500) was added to each metabolite sample and 400 microliters of 2:1 chloroform:methanol (OmniSolv® Chloroform, Sigma Aldrich, CAT CX1059) was added to each lipid sample. All samples were kept on dry ice during shipping then stored at -80°C upon arrival at Biology Centre, České Budějovice until processing.

Each sample was thawed on ice and subjected to homogenization using a DISPO X120 CAT metal blade-homogenizer starting at the lowest rotation speed then gradually increasing to maximum for a duration of ca. 10 seconds at room temperature. Blades were cleaned between samples using a chloroform:methanol solution. Homogenized samples were put on ice for 60 minutes to allow time for extraction, with vortexing every 10 minutes, before being centrifuged at 20,000 g at 4°C for either ten minutes (for metabolite samples) or two minutes (for lipid samples). For each metabolite sample, 300-μL of supernatant was collected and stored at -80°C until analysis. For each lipid sample, 300-μL of supernatant was collected, combined with 60-μL of 0.9% NaCl, mixed, vortexed, then centrifuged again at 20,000 g for 2 minutes at 4°C. The organic chloroform phase of each lipid sample (ca. 200 microliters) was removed, placed into a glass vial with a Teflon lid, dried under nitrogen, then stored at -80°C until analysis.

*4.2 LC/MS metabolite analysis*

We utilized the Q Exactive Plus Orbitrap mass spectrometer coupled to a Dionex Ultimate 3000 liquid chromatograph and a Dionex open autosampler (all from ThermoFisher Scientific, Waltham, MA, USA). Briefly, separate runs were conducted with a mass range of 70 to 1000 Da at 70,000 resolution (at mass m/z 200) to record full scan LC/MS positive (PESI) and negative (NESI) ion mass spectra. The settings of the LC-MS were: scan rate ±3 Hz, 3×106 automatic gain control (AGC) target, and maximum ion injection time (IT) 100 ms. Source ionization parameters were as follows: ±3000 V spray voltage, 350°C capillary temperature, sheath gas at 60 au, aux gas at 20 au, spare gas at 1 au, probe temperature 350°C and S-Lens level at 60 au. For accurate mass identification, we used lock masses of 622.0290 Da for the positive ion mode and 301.9981 Da for the negative ion mode. A SeQuant ZIC-pHILIC HPLC column (150 mm, 4.6 mm i.d., 5 m, Merck, Darmstadt, Germany) was used for chromatographic separation of metabolites with the following parameters: mobile phase flow rate 450 l min−1, injection volume 5 l, column temperature 35°C; mobile phase: A=acetonitrile, B=20 mmol l−1 aqueous ammonium carbonate (pH 9.2 adjusted by NH4OH, Sigma-Aldrich); gradient: 0 min, 20% B; 20 min, 80% B; 20.1 min, 95% B; 23.3 min, 95% B; 23.4 min, 20% B; 30.0 min 20% B. Quality control samples (extracts pooled from all samples) and blank samples (no eggs present) were included periodically in the analysis. We then used an in-house Metabolite Mapper platform equipped with an internal metabolite database in conjunction with Xcalibur™ software (v.2.1, ThermoFisher Scientific) to acquire data and identify metabolites.

#### *4.3 Lipid extraction*

Total lipids were extracted using a simplified Folch method (Folch et al., 1957). Each sample was homogenized in 1.5 mL tubes containing 300 µL ice-cold chloroform:methanol (2:1 v/v; Honeywell International, Prague, Czech Republic) using a pellet pestle and battery-driven homogenizer. Samples were centrifuged for 10 min at 20,000 g, 4°C, and the supernatant transferred to a clean tube. Pellets were re-extracted with 300 µL solvent, and supernatants were pooled. Phase separation was induced by adding 120 µL 0.9% NaCl, vortexing, and centrifugation for 10 min at 20,000 g, 4°C (Universal 32R). The lower organic phase was collected into 2 mL glass vials with PTFE-lined caps, evaporated under nitrogen, and stored at -80°C until further processing. Non-polar lipids (triacylglycerols) were removed by re-dissolving the dry lipids in 600 µL hexane and 600 µL 80% acetonitrile in water, vortexing, ultrasonic treatment for 10 min, and centrifugation for 5 min at 700 g. The lower acetonitrile phase containing polar lipids (mainly phospholipids) was collected, evaporated, and stored at -80°C. Each sample was split into two identical aliquots for LC/HRMS analysis of phospholipid (PL) classes and GC/MS analysis of fatty acid methyl esters (FAMEs).

*4.4 LC/HRMS analysis of PL classes*

Dry lipid samples were dissolved in 60 µL methanol:isopropanol (1:1 v/v; Honeywell). PL classes were separated and identified by liquid chromatography coupled to high-resolution mass spectrometry (LC/HRMS) using an Orbitrap Q Exactive Plus coupled to a Dionex Ultimate 3000 LC and open autosampler (Thermo Fisher Scientific, San Jose, CA, USA). The method was a slight modification of (Cífková et al., 2012). Chromatographic conditions: injection volume 5 µL; Kinetex 2.6 µm HILIC column (100 Å, 150 × 2.1 mm I.D., Phenomenex, Torrance, CA, USA) at 35°C; mobile phases: (A) acetonitrile and (B) 5 mM ammonium acetate in water. Gradient: 0 min: 94% A; 34 min: 84% A; 40 min: 77% A; 40.1 min: 94% A; 60 min: 94% A; flow rate 350 µL/min. Electrospray ionization: positive ion mode (PESI, –3.3 kV); capillary temp 250°C; sheath gas 30 au; auxiliary gas 10 au; sweep gas 1 au; S-lens RF 60; auxiliary gas heater 350°C. Full scan m/z 300–1600 at 70,000 resolution (m/z 200); scan rate ±3 Hz; AGC target $3 \times 10^6$; max IT 100 ms. PL identification was performed using data-dependent MS/MS (dd-$MS^2$) at 17,500 resolution, AGC target $1 \times 10^5$, max IT 50 ms, isolation window 4 m/z, NCE 30. Accurate mass measurements used natural lock masses: m/z 391.2843 and 279.1511. Data acquisition and processing were performed in XCalibur 4.0. Peak areas of all lipids in each PL class were summed, and relative abundance expressed as a percentage of the total.

*4.5 GC/MS analysis of fatty acids*

FAMEs were analyzed using a 7890B GC coupled to 7010B triple quadrupole MS (Agilent, Santa Clara, CA, USA), following a slightly modified method from (Vodrážka et al., 2025). Heptadecanoic-$d_{33}$ acid (5 µL, 0.1 nmol/µL; Merck) was added as an internal standard, and samples were transesterified with sodium methoxide (Merck). Specifically, 50 µL methyl tert-butyl ether and 100 µL 2 M sodium methoxide in methanol were added, vortexed 3 min, followed by 100 µL hexane and 100 µL 2 M HCl, mixed, and phase-separated. Upper hexane layers were collected, aqueous phases re-extracted, combined, evaporated, and dissolved in 50 µL isooctane. Aliquots (1 µL) were injected onto a 30 m ZB-FAME column (0.25 mm I.D., 0.20 µm film; Phenomenex). GC parameters: He flow 1.1 mL/min; inlet 250°C; splitless injection 0.5 min; oven 70°C 2 min → 15°C/min to 180°C hold 6 min → 10°C/min to 230°C hold 1 min; transfer line 250°C; ion source 230°C; EI 70 eV. Data were acquired and processed in Agilent MassHunter v10. Absolute quantification used Supelco 37 Component FAME Mix (CRM 47885) as external standard; heptadecanoic-$d_{33}$ acid corrected for preparation and injection variability. FAMEs were expressed as molar percentages of total FAME content.

### 5. Statistical Analyses

To account for slight differences in control (i.e., non-cold-shocked) egg viability across treatment groups, prior to modeling cold shock survival after exposure to -9°C and -11°C, we scaled raw survival (i.e., number of hatched larvae / number of eggs) of each replicate paper to the average survival of the control group exposed to the same diapause and acclimation treatment combination (i.e., divided survival of each replicate by average survival of the appropriate control group). Thus, survival data was modeled as a proportion, with the number of binomial trials (i.e., total egg count) per replicate specified using the weights parameter. Because the cold shock survival of individual replicates sometimes exceeded the average survival of the relevant control group, the scaled survival value in some cases slightly exceeded one (max: 1.04). These values were retained in the final datasets and figures, but for the purposes of statistical analyses, all values were bounded by [0,1] to allow for modeling with beta-binomial distribution. In the -11°C survival dataset, all replicates in the non-diapause before acclimation group exhibited zero survival, preventing proper convergence of the initial generalized linear mixed effects model due to complete separation. To overcome this issue, we ran a model with the same structure (fixed effects of diapause treatment, acclimation treatment, days post-oviposition, and the interaction between diapause and acclimation treatments), but which excluded replicates from the non-diapause before acclimation group and the diapause before acclimation group. We used this model to report results related to the significance of our fixed effects (i.e., to determine whether diapause, acclimation treatment, and their interaction affected cold shock survival). To perform post-hoc comparisons of individual treatment groups using the emmeans function, we ran a new model with a fixed effect of total treatment (i.e., the combination of diapause and acclimation treatments) and days post-oviposition. This new model was run using all replicates except those from the non-diapause before acclimation treatment group.

For all linear mixed effects models of individuals metabolites, PL classes, and FAME species, data was either log or square-root transformed to improve normality of residuals (assessed using a Shapiro-Wilk normality test via shapiro.test function in R stats package). If the data failed to meet the assumption of normality of residuals after transformations (i.e., a $p < 0.05$ in Shapiro-Wilk normality test), we instead tested for a significant effect of total treatment using a non-parametric Kruskal-Wallis test (kruskal.test function in R stats package), followed by the same planned set of five post-hoc pairwise contrasts using non-parametric Wilcoxon rank sum tests (wilcox.test function in R stats package). *P*-values generated for each pairwise contrast were adjusted to account for multiple testing across the five pairwise contrasts and the total number of response variables tested per dataset (Benjamini & Hochberg, 1995).

### 6. Supplementary Information References

Batz, Z. A., Clemento, A. J., Fritzenwanker, J., Ring, T. J., Garza, J. C., & Armbruster, P. A. (2020). Rapid adaptive evolution of the diapause program during range expansion of an invasive mosquito. *Evolution*, *74*(7), 1451–1465. https://doi.org/10.1111/evo.14029

Benjamini, Y., & Hochberg, Y. (1995). Controlling the false discovery rate: a practical and powerful approach to multiple testing. *Journal of the Royal Statistical Society. Series B (Methodological)*, *57*(1), 289–300. https://doi.org/10.1111/j.2517-6161.1995.tb02031.x

Cífková, E., Holčapek, M., Lísa, M., Ovčačíková, M., Lyčka, A., Lynen, F., & Sandra, P. (2012). Nontargeted quantitation of lipid classes using hydrophilic interaction liquid chromatography-electrospray ionization mass spectrometry with single internal standard and response factor approach. *Analytical Chemistry*, *84*(22), 10064–10070. https://doi.org/10.1021/ac3024476

Folch, J., Lees, M., & Sloane Stanley, G. H. (1957). A simple method for the isolation and purification of total lipides from animal tissues. *Journal of Biological Chemistry*, *226*(1), 497–509.

Hanson, S. M., & Craig, G. B. (1994). Cold acclimation, diapause, and geographic origin affect cold hardiness in eggs of Aedes albopictus (Diptera: Culicidae). *Journal of Medical Entomology*, *31*(2), 192–201. https://doi.org/10.1093/jmedent/31.2.192

Hanson, S. M., & Craig, G. B. (1995). Relationship between cold hardiness and supercooling point in Aedes albopictus eggs. *Journal of the American Mosquito Control Association*, *11*(1), 35–38.

Trpiš, M. (1970). A new bleaching and decalcifying method for general use in zoology. *Canadian Journal of Zoology*, *48*(4), 892–893.

Vodrážka, P., Řimnáčová, L., Berková, P., Vojtíšek, J., Verner, M., Moos, M., & Šimek, P. (2025). A sew, validated GC-PICI-MS method for the quantification of 32 lipid fatty acids via base-catalyzed transmethylation and the isotope-coded derivatization of internal standards. *Metabolites*, *15*(2), 104. https://doi.org/10.3390/metabo15020104

### Supplementary Figures

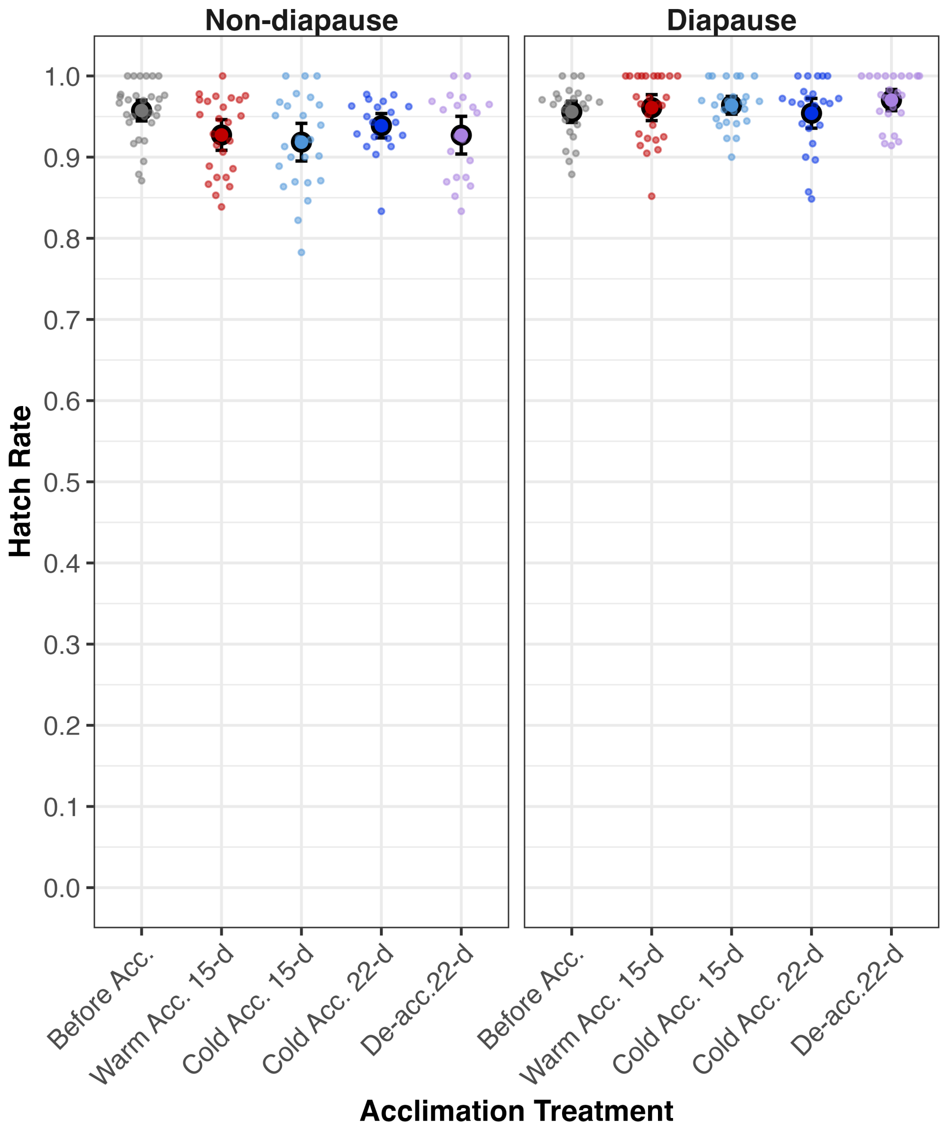


**Figure S1** Control (non-cold-shocked) survival (i.e., hatch rates) of each treatment group. Large points and error bars indicate the mean survival rates ± two standard errors of each treatment group, while smaller points show the proportion of surviving individuals in each replicate. Colors are as in Figure 1.

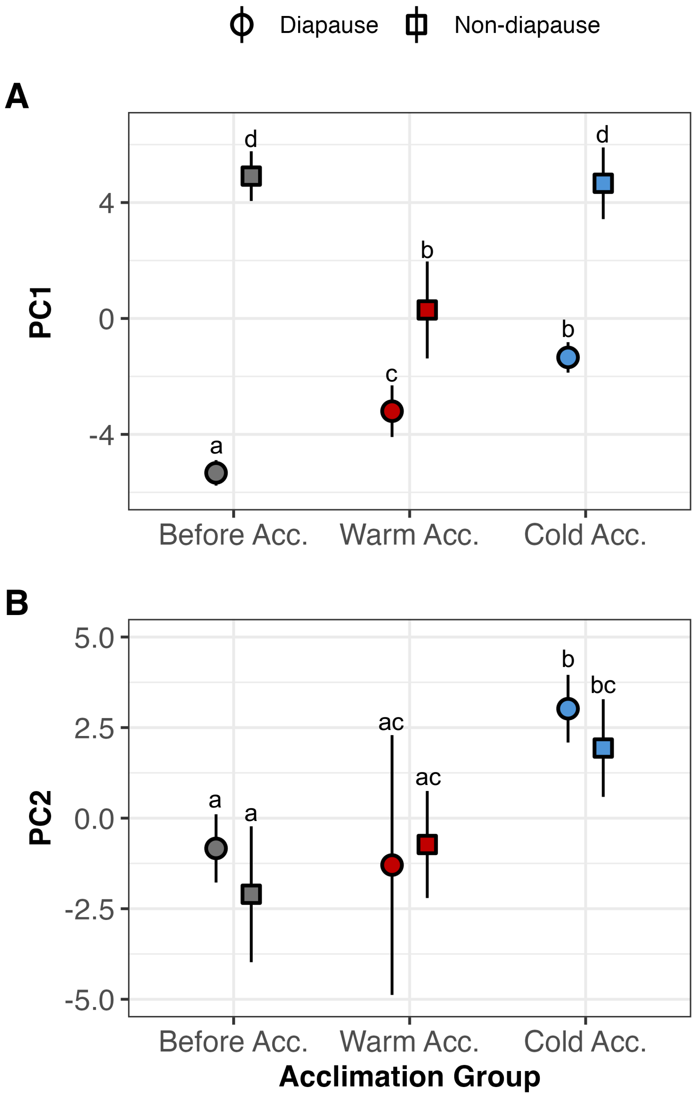


**Figure S2** Values of the first **(A)** and second **(B)** principal components (PCs) for the metabolite profile PC analysis. Points and error bars indicate the mean PC values ± two standard errors of each treatment group. Points with different letters are significantly different (p-adjusted < 0.05) in post-hoc testing. Treatment abbreviations are as follows: Before Acc. = before acclimation; Warm Acc. = warm acclimation 15-d; Cold Acc. = cold acclimation 15-d).

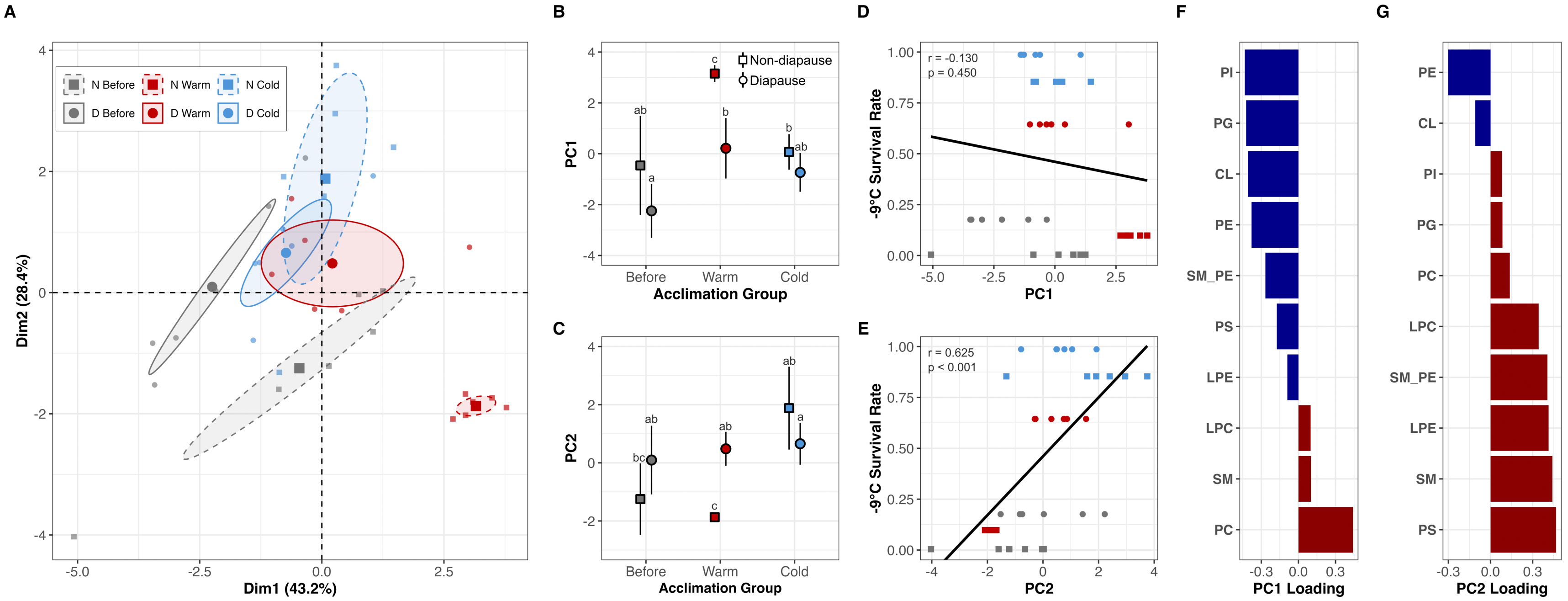


**Figure S3** Phospholipid (PL) class profile is affected by both diapause and cold acclimation. **(A)** Principal component analysis (PCA) plot showing separation of egg samples based on variation in PL class profiles. Each individual sample is shown as a small point, with the group mean and 95% confidence interval represented by a larger point surrounded by a shaded ellipse. Values of the first **(B)** and second **(C)** PCs from the PCA shown in **(A)**. Points and error bars indicate the mean survival rates ± two standard errors of each treatment group. Points with different letters are significantly different (p- adjusted < 0.05) in post-hoc testing. Correlations (Pearson coefficient) between the first two PC dimensions and the average -9°C cold shock survival of each treatment group are shown in **(D)** and **(E)**, with color and shape formatting as indicated by the legend in **(A)**. PL classes contributing to variation in PC1 **(D)** and PC2 **(E)** ranked according to their loading value (positive values as red and negative values as blue). Treatment abbreviations are as follows: ND = non-diapause; D = diapause; Before = before acclimation; Warm = warm acclimation 15-d; Cold = cold acclimation 15-d).

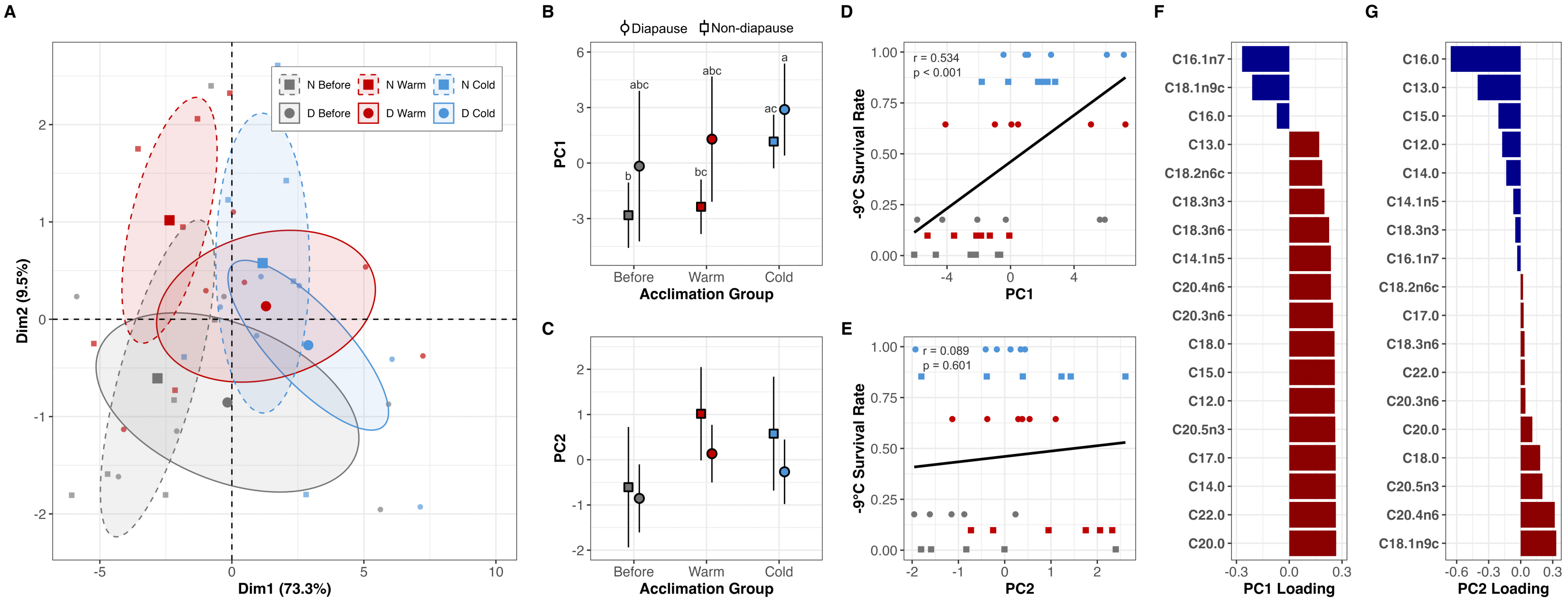


**Figure S4** Fatty acid methyl ester (FAME) profile principal component analysis (PCA) results. **(A)** PCA plot showing separation of egg samples based on variation in FAME profiles. Each individual sample is shown as a small point, with the group mean and 95% confidence interval represented by a larger point surrounded by a shaded ellipse. Values of the first **(B)** and second **(C)** PCs from the PCA shown in **(A)**. Points and error bars indicate the mean survival rates ± two standard errors of each treatment group. Points with different letters are significantly different (p- adjusted < 0.05) in post-hoc testing. Correlations (Pearson coefficient) between the first two PC dimensions and the average -9°C cold shock survival of each treatment group are shown in **(D)** and **(E)**, with color and shape formatting as indicated by the legend in **(A)**. FAME species contributing to variation in PC1 **(D)** and PC2 **(E)** ranked according to their loading value (positive values as red and negative values as blue). Treatment abbreviations are as follows: ND = non-diapause; D = diapause; Before = before acclimation; Warm = warm acclimation 15-d; Cold = cold acclimation 15-d).

### Supplementary Tables

**Table S1** Replication statement for cold shock survival trials and biochemical analyses.

| **Experiment** | **Scale of inference** | **Scale at which the factor of interest is applied** | **Number of replicates at the appropriate scale** |
|---|---|---|---|
| Cold Shock Survival | egg paper replicate (groups of on average 36 eggs) | egg paper replicate | 20-31 egg paper replicates for each of 30 total treatment combinations |
| Biochemical Analyses | egg paper replicate (average of 6.13 mg of eggs) | egg paper replicate | 6 egg paper replicates for each of 6 total treatment combinations for each assay (metabolite and lipid) |

**Table S2** List of which combinations of diapause, acclimation, and cold shock treatments each replicate parental cage contributed eggs to in cold shock experiment. Treatment abbreviations are as follows: ND = non-diapause; D = diapause; Before = before acclimation; Warm = warm acclimation; Cold = cold acclimation; Deacc = de-acclimation.

| Parental Cage ID | Number of Total Treatments | Total Treatment List |
|---|---|---|
| 1 | 9 | ND Before -11°C, ND Before -9°C, ND Before Control, ND Warm 15-d -11°C, ND Warm 15-d -9°C, ND Warm 15-d Control, ND Cold 15-d -11°C, ND Cold 15-d -9°C, ND Cold 15-d Control |
| 2 | 8 | ND Before -9°C, ND Before Control, ND Warm 15-d -11°C, ND Warm 15-d -9°C, ND Warm 15-d Control, ND Cold 15-d -11°C, ND Cold 15-d -9°C, ND Cold 15-d Control |
| 3 | 10 | D Before -11°C, D Before -9°C, D Before Control, D Warm 15-d -11°C, D Warm 15-d -9°C, D Warm 15-d Control, D Cold 15-d -11°C, D Cold 15-d -9°C, D Cold 15-d Control, D Cold 22-d Control |
| 4 | 9 | D Before -11°C, D Before -9°C, D Before Control, D Warm 15-d -11°C, D Warm 15-d -9°C, D Warm 15-d Control, D Cold 15-d -11°C, D Cold 15-d -9°C, D Cold 15-d Control |
| 5 | 9 | D Before -11°C, D Before -9°C, D Before Control, D Warm 15-d -11°C, D Warm 15-d -9°C, D Warm 15-d Control, D Cold 15-d -11°C, D Cold 15-d -9°C, D Cold 15-d Control |
| 6 | 15 | D Before -11°C, D Before -9°C, D Before Control, D Warm 15-d -11°C, D Warm 15-d -9°C, D Warm 15-d Control, D Cold 15-d -11°C, D Cold 15-d -9°C, D Cold 15-d Control, D Cold 22-d -11°C, D Cold 22-d -9°C, D Cold 22-d Control, D Deacc 22-d -11°C, D Deacc 22-d -9°C, D Deacc 22-d Control |
| 7 | 15 | ND Before -11°C, ND Before -9°C, ND Before Control, ND Warm 15-d -11°C, ND Warm 15-d -9°C, ND Warm 15-d Control, ND Cold 15-d -11°C, ND Cold 15-d -9°C, ND Cold 15-d Control, ND Cold-22d -11°C, ND Cold-22d -9°C, ND Cold 22-d Control, ND Deacc 22-d -11°C, ND Deacc 22-d -9°C, ND Deacc 22-d Control |
| 8 | 9 | ND Before -11°C, ND Before -9°C, ND Before Control, ND Warm 15-d -11°C, ND Warm 15-d -9°C, ND Warm 15-d Control, ND Cold 15-d -11°C, ND Cold 15-d -9°C, ND Cold 15-d Control |
| 9 | 10 | D Warm 15-d -9°C, D Warm 15-d Control, D Cold 15-d -9°C, D Cold 15-d Control, D Cold 22-d -11°C, D Cold 22-d -9°C, D Cold 22-d Control, D Deacc 22-d -11°C, D Deacc 22-d -9°C, D Deacc 22-d Control |
| 10 | 6 | D Cold 22-d -11°C, D Cold 22-d -9°C, D Cold 22-d Control, D Deacc 22-d -11°C, D Deacc 22-d -9°C, D Deacc 22-d Control |
| 11 | 6 | ND Cold 22-d -11°C, ND Cold 22-d -9°C, ND Cold 22-d Control, ND Deacc 22-d -11°C, ND Deacc 22-d -9°C, ND Deacc 22-d Control |
| 12 | 6 | D Cold 22-d -11°C, D Cold 22-d -9°C, D Cold 22-d Control, D Deacc 22-d -11°C, D Deacc 22-d -9°C, D Deacc 22-d Control |
| 13 | 15 | ND Before -11°C, ND Before -9°C, ND Before Control, ND Warm 15-d -11°C, ND Warm 15-d -9°C, ND Warm 15-d Control, ND Cold 15-d -11°C, ND Cold 15-d -9°C, ND Cold 15-d Control, ND Cold-22d -11°C, ND Cold 22-d -9°C, ND Cold 22-d Control, ND Deacc 22-d -11°C, ND Deacc 22-d -9°C, ND Deacc 22-d Control |
| 14 | 6 | ND Cold 22-d -11°C, ND Cold 22-d -9°C, ND Cold 22-d Control, ND Deacc 22-d -11°C, ND Deacc 22-d -9°C, ND Deacc 22-d Control |

**Table S3** The number of individual egg paper replicates used for each combination of diapause, acclimation, and cold-shock treatments. Each egg paper replicate contained an average of 36 eggs.

| Diapause Treatment | Acclimation Treatment | Cold Shock Treatment | N |
|---|---|---|---|
| Non-diapause | Before Acclimation | -11°C for 20hrs | 28 |
| Non-diapause | Before Acclimation | -9°C for 20hrs | 31 |
| Non-diapause | Before Acclimation | Control | 30 |
| Non-diapause | Warm Acclimated 15-d | -11°C for 20hrs | 25 |
| Non-diapause | Warm Acclimated 15-d | -9°C for 20hrs | 28 |
| Non-diapause | Warm Acclimated 15-d | Control | 25 |
| Non-diapause | Cold Acclimated 15-d | -11°C for 20hrs | 25 |
| Non-diapause | Cold Acclimated 15-d | -9°C for 20hrs | 28 |
| Non-diapause | Cold Acclimated 15-d | Control | 25 |
| Non-diapause | Cold Acclimated 22-d | -11°C for 20hrs | 21 |
| Non-diapause | Cold Acclimated 22-d | -9°C for 20hrs | 20 |
| Non-diapause | Cold Acclimated 22-d | Control | 20 |
| Non-diapause | De-acclimated 22-d | -11°C for 20hrs | 21 |
| Non-diapause | De-acclimated 22-d | -9°C for 20hrs | 20 |
| Non-diapause | De-acclimated 22-d | Control | 20 |
| Diapause | Before Acclimation | -11°C for 20hrs | 26 |
| Diapause | Before Acclimation | -9°C for 20hrs | 28 |
| Diapause | Before Acclimation | Control | 25 |
| Diapause | Warm Acclimated 15-d | -11°C for 20hrs | 25 |
| Diapause | Warm Acclimated 15-d | -9°C for 20hrs | 26 |
| Diapause | Warm Acclimated 15-d | Control | 26 |
| Diapause | Cold Acclimated 15-d | -11°C for 20hrs | 25 |
| Diapause | Cold Acclimated 15-d | -9°C for 20hrs | 26 |
| Diapause | Cold Acclimated 15-d | Control | 25 |
| Diapause | Cold Acclimated 22-d | -11°C for 20hrs | 21 |
| Diapause | Cold Acclimated 22-d | -9°C for 20hrs | 20 |
| Diapause | Cold Acclimated 22-d | Control | 23 |
| Diapause | De-acclimated 22-d | -11°C for 20hrs | 21 |
| Diapause | De-acclimated 22-d | -9°C for 20hrs | 20 |
| Diapause | De-acclimated 22-d | Control | 23 |

**Table S4** List of the 50 metabolites detected in egg samples, with abbreviated names used in main text figures.

| Full Name | Abbreviation |
|---|---|
| 2 and 3-phosphoglyceric acid | 2 and 3-phosphoglyceric acid |
| 2-Oxoglutaric acid | 2-Oxoglutaric acid |
| Acetylcholine | A-Chol |
| ADP | ADP |
| Alanine | Ala |
| AMP | AMP |
| Arachidic acid | Arachidic acid |
| Arginine | Arg |
| Asparagine | Asn |
| Aspartic acid | Aspartic acid |
| ATP | ATP |
| Betaine | Betaine |
| CDP-choline | CDP-Chol |
| Citric acid | Citric acid |
| Citrulline | Cit |
| Cystathionine | Cystathionine |
| D and L Glutamic acid | Glutamic acid |
| D and L Lactic acid | Lactic acid |
| Fructose-1P and 6P and Glucose-1P | Fructose-1P and 6P and Glucose-1P |
| Fumaric acid | Fumaric acid |
| Glucose | Glucose |
| Glucose-6P | Glucose-6P |
| Glutamine | Gln |
| Glyceric acid | Glyceric acid |
| Glycerol | Glycerol |
| Glycerophosphoethanolamine | GP-Etn |
| Glycerophosphocholine | GP-Chol |
| Choline | Chol |
| Leucine | Leu |
| Linoleic acid | Linoleic acid |
| Malic acid | Malic acid |
| Methionine | Met |
| Oleic acid | Oleic acid |
| Palmitic acid | Palmitic acid |
| Palmitoleic acid | Palmitoleic acid |
| Phenylalanine | Phe |

| | |
|---|---|
| Phosphoarginine | P-Arg |
| Phosphocholine | P-Chol |
| Proline | Pro |
| Pyruvic acid | Pyruvic acid |
| Ribitol Adonitol | Ribitol |
| Serine | Ser |
| Sorbitol | Sorbitol |
| Stearic acid | Stearic acid |
| Succinic acid | Succinic acid |
| Threonine | Thr |
| Trehalose | Trehalose |
| Tryptophan | Trp |
| Tyrosine | Tyr |
| Valine | Val |

**Table S5** Results of post-hoc testing comparing cold shock survival after exposure to -9°C for 20 hours in eggs of different diapause and acclimation treatments. Treatment abbreviations are as follows: ND = non-diapause; D = diapause; Before = before acclimation; Warm = warm acclimation; Cold = cold acclimation; DeAcc = de-acclimation. Significant contrasts are bolded.

| Contrast | Estimate | SE | z-ratio | *p*-value |
|---|---|---|---|---|
| **ND Before – D Before** | -3.58 | 0.53 | -6.81 | 1.39E-11 |
| **ND Before – ND Warm 15-d** | -2.97 | 0.52 | -5.66 | 1.89E-08 |
| **ND Before – D Warm 15-d** | -5.74 | 0.53 | -10.87 | 2.97E-27 |
| **ND Before – ND Cold 15-d** | -6.92 | 0.52 | -13.28 | 6.37E-40 |
| **ND Before – D Cold 15-d** | -8.82 | 0.59 | -14.95 | 5.98E-50 |
| **ND Before – ND Cold 22-d** | -7.87 | 0.55 | -14.35 | 3.02E-46 |
| **ND Before – D Cold 22-d** | -9.40 | 0.64 | -14.60 | 9.80E-48 |
| **ND Before – ND DeAcc 22-d** | -3.89 | 0.52 | -7.46 | 1.34E-13 |
| **ND Before – D DeAcc 22-d** | -6.31 | 0.53 | -11.83 | 5.27E-32 |
| **D Before – ND Warm 15-d** | 0.61 | 0.21 | 2.96 | 3.29E-03 |
| **D Before – D Warm 15-d** | -2.16 | 0.16 | -13.57 | 1.48E-41 |
| **D Before – ND Cold 15-d** | -3.34 | 0.19 | -17.35 | 2.22E-66 |
| **D Before – D Cold 15-d** | -5.24 | 0.31 | -17.08 | 1.91E-64 |
| **D Before – ND Cold 22-d** | -4.29 | 0.25 | -16.99 | 6.02E-64 |
| **D Before – D Cold 22-d** | -5.82 | 0.40 | -14.46 | 7.38E-47 |
| D Before – ND DeAcc 22-d | -0.31 | 0.19 | -1.66 | 9.98E-02 |
| **D Before – D DeAcc 22-d** | -2.73 | 0.19 | -14.30 | 5.97E-46 |
| **ND Warm 15-d – D Warm 15-d** | -2.77 | 0.20 | -14.18 | 3.40E-45 |
| **ND Warm 15-d – ND Cold 15-d** | -3.95 | 0.18 | -21.47 | 1.43E-100 |
| **ND Warm 15-d – D Cold 15-d** | -5.85 | 0.33 | -18.00 | 3.17E-71 |
| **ND Warm 15-d – ND Cold 22-d** | -4.90 | 0.26 | -19.15 | 2.13E-80 |
| **ND Warm 15-d – D Cold 22-d** | -6.43 | 0.41 | -15.65 | 1.61E-54 |
| **ND Warm 15-d – ND DeAcc 22-d** | -0.92 | 0.20 | -4.69 | 3.29E-06 |
| **ND Warm 15-d – D DeAcc 22-d** | -3.34 | 0.21 | -15.97 | 1.02E-56 |
| **D Warm 15-d – ND Cold 15-d** | -1.17 | 0.18 | -6.58 | 6.59E-11 |
| **D Warm 15-d – D Cold 15-d** | -3.08 | 0.29 | -10.66 | 2.93E-26 |
| **D Warm 15-d – ND Cold 22-d** | -2.13 | 0.25 | -8.63 | 1.05E-17 |
| **D Warm 15-d – D Cold 22-d** | -3.65 | 0.39 | -9.44 | 6.56E-21 |
| **D Warm 15-d – ND DeAcc 22-d** | 1.85 | 0.18 | 10.10 | 9.56E-24 |
| **D Warm 15-d – D DeAcc 22-d** | -0.57 | 0.16 | -3.47 | 5.78E-04 |
| **ND Cold 15-d – D Cold 15-d** | -1.90 | 0.31 | -6.06 | 1.76E-09 |
| **ND Cold 15-d – ND Cold 22-d** | -0.96 | 0.24 | -3.93 | 9.98E-05 |
| **ND Cold 15-d – D Cold 22-d** | -2.48 | 0.40 | -6.18 | 8.60E-10 |
| **ND Cold 15-d – ND DeAcc 22-d** | 3.02 | 0.18 | 16.67 | 1.29E-61 |
| **ND Cold 15-d – D DeAcc 22-d** | 0.61 | 0.19 | 3.16 | 1.74E-03 |

| | | | | |
|---|---|---|---|---|
| **D Cold 15-d – ND Cold 22-d** | 0.95 | 0.36 | 2.65 | 8.43E-03 |
| D Cold 15-d – D Cold 22-d | -0.57 | 0.46 | -1.25 | 2.13E-01 |
| **D Cold 15-d – ND DeAcc 22-d** | 4.93 | 0.32 | 15.47 | 2.18E-53 |
| **D Cold 15-d – D DeAcc 22-d** | 2.51 | 0.30 | 8.26 | 2.25E-16 |
| **ND Cold 22-d – D Cold 22-d** | -1.52 | 0.44 | -3.50 | 5.44E-04 |
| **ND Cold 22-d – ND DeAcc 22-d** | 3.98 | 0.23 | 17.06 | 2.31E-64 |
| **ND Cold 22-d – D DeAcc 22-d** | 1.56 | 0.26 | 6.08 | 1.61E-09 |
| **D Cold 22-d – ND DeAcc 22-d** | 5.50 | 0.41 | 13.54 | 2.16E-41 |
| **D Cold 22-d – D DeAcc 22-d** | 3.09 | 0.38 | 8.02 | 1.58E-15 |
| **ND DeAcc 22-d – D DeAcc 22-d** | -2.42 | 0.20 | -12.12 | 1.83E-33 |

**Table S6** Results of post-hoc testing comparing cold shock survival after exposure to -11°C for 20 hours in eggs of different diapause and acclimation treatments. Treatment abbreviations are as follows: ND = non-diapause; D = diapause; Before = before acclimation; Warm = warm acclimation; Cold = cold acclimation; DeAcc = de-acclimation. Significant contrasts are bolded.

| Contrast | Estimate | SE | z-ratio | *p*-value |
|---|---|---|---|---|
| **D Before – D Cold 15-d** | -6.89 | 0.49 | -13.97 | 1.16E-43 |
| **D Before – D Cold 22-d** | -7.05 | 0.52 | -13.67 | 7.30E-42 |
| **D Before – D DeAcc 22-d** | -2.09 | 0.53 | -3.99 | 9.62E-05 |
| **D Before – D Warm 15-d** | -1.31 | 0.53 | -2.46 | 1.67E-02 |
| **D Before – ND Cold 15-d** | -4.26 | 0.50 | -8.48 | 4.03E-17 |
| **D Before – ND Cold 22-d** | -4.87 | 0.51 | -9.64 | 1.17E-21 |
| D Before – ND DeAcc 22-d | -0.60 | 0.60 | -1.00 | 3.25E-01 |
| D Before – ND Warm 15-d | 1.57 | 1.12 | 1.40 | 1.70E-01 |
| D Cold 15-d – D Cold 22-d | -0.16 | 0.24 | -0.68 | 4.95E-01 |
| **D Cold 15-d – D DeAcc 22-d** | 4.79 | 0.26 | 18.40 | 1.68E-74 |
| **D Cold 15-d – D Warm 15-d** | 5.58 | 0.27 | 20.44 | 1.35E-91 |
| **D Cold 15-d – ND Cold 15-d** | 2.63 | 0.20 | 12.91 | 1.65E-37 |
| **D Cold 15-d – ND Cold 22-d** | 2.01 | 0.22 | 9.26 | 3.92E-20 |
| **D Cold 15-d – ND DeAcc 22-d** | 6.29 | 0.39 | 16.28 | 8.61E-59 |
| **D Cold 15-d – ND Warm 15-d** | 8.45 | 1.02 | 8.29 | 1.82E-16 |
| **D Cold 22-d – D DeAcc 22-d** | 4.95 | 0.24 | 20.59 | 1.24E-92 |
| **D Cold 22-d – D Warm 15-d** | 5.74 | 0.31 | 18.31 | 6.06E-74 |
| **D Cold 22-d – ND Cold 15-d** | 2.79 | 0.23 | 11.88 | 4.65E-32 |
| **D Cold 22-d – ND Cold 22-d** | 2.18 | 0.23 | 9.30 | 2.94E-20 |
| **D Cold 22-d – ND DeAcc 22-d** | 6.45 | 0.40 | 16.32 | 4.76E-59 |
| **D Cold 22-d – ND Warm 15-d** | 8.62 | 1.03 | 8.40 | 7.63E-17 |
| **D DeAcc 22-d – D Warm 15-d** | 0.79 | 0.32 | 2.42 | 1.81E-02 |
| **D DeAcc 22-d – ND Cold 15-d** | -2.16 | 0.25 | -8.58 | 1.82E-17 |
| **D DeAcc 22-d – ND Cold 22-d** | -2.78 | 0.25 | -11.12 | 2.65E-28 |
| **D DeAcc 22-d – ND DeAcc 22-d** | 1.50 | 0.40 | 3.73 | 2.64E-04 |
| **D DeAcc 22-d – ND Warm 15-d** | 3.66 | 1.03 | 3.56 | 4.90E-04 |
| **D Warm 15-d – ND Cold 15-d** | -2.95 | 0.29 | -10.16 | 7.76E-24 |
| **D Warm 15-d – ND Cold 22-d** | -3.57 | 0.29 | -12.12 | 2.70E-33 |
| D Warm 15-d – ND DeAcc 22-d | 0.71 | 0.43 | 1.65 | 1.08E-01 |
| **D Warm 15-d – ND Warm 15-d** | 2.88 | 1.04 | 2.77 | 6.93E-03 |
| **ND Cold 15-d – ND Cold 22-d** | -0.62 | 0.19 | -3.24 | 1.56E-03 |
| **ND Cold 15-d – ND DeAcc 22-d** | 3.66 | 0.37 | 9.89 | 1.14E-22 |
| **ND Cold 15-d – ND Warm 15-d** | 5.83 | 1.01 | 5.79 | 1.06E-08 |
| **ND Cold 22-d – ND DeAcc 22-d** | 4.27 | 0.35 | 12.36 | 1.58E-34 |

| | | | | |
|---|---|---|---|---|
| **ND Cold 22-d – ND Warm 15-d** | 6.44 | 1.02 | 6.34 | 3.65E-10 |
| **ND DeAcc 22-d – ND Warm 15-d** | 2.17 | 1.06 | 2.04 | 4.63E-02 |

**Table S7** The relative proportions of each phospholipid class found in each treatment group. Values are grouped means (averaged across the six replicates of each treatment group) +/- two standard errors.

| Abbreviation | Full Name | ND Before | ND Warm 15-d | ND Cold 15-d | D Before | D Warm 15-d | D Cold 15-d |
|---|---|---|---|---|---|---|---|
| PC | phosphatidylcholine | 46.35%±7.13 | 57.03%±1.69 | 49.7%±3.68 | 44.68%±5.06 | 50.3%±3.6 | 45.93%±3.12 |
| PE | phosphoethanolamine | 44.02%±7.84 | 35.15%±1.72 | 37.73%±4.72 | 43.2%±5.93 | 37.88%±2.79 | 41.52%±3.55 |
| SM_PE | ethanolamine sphingomyelin | 4.32%±0.36 | 3.2%±0.28 | 5.09%±0.26 | 5.11%±0.33 | 4.3%±0.81 | 5.1%±0.21 |
| SM | sphingomyelin[1] | 2.65%±0.6 | 2.73%±0.31 | 3.52%±0.5 | 2.87%±0.53 | 4.12%±0.32 | 3.67%±0.37 |
| PS | phosphatidylserine | 1.2%±0.26 | 1.23%±0.12 | 2.22%±0.37 | 2.19%±0.24 | 1.88%±0.37 | 2.26%±0.23 |
| PI | phosphatidylinositol | 0.71%±0.21 | 0.27%±0.04 | 0.7%±0.14 | 1.24%±0.16 | 0.85%±0.1 | 0.78%±0.12 |
| LPE | lysophosphatidylethanolamine | 0.48%±0.07 | 0.27%±0.04 | 0.76%±0.15 | 0.44%±0.08 | 0.4%±0.06 | 0.48%±0.09 |
| CL | cardiolipin | 0.12%±0.09 | 0.04%±0.01 | 0.1%±0.02 | 0.14%±0.02 | 0.11%±0.02 | 0.11%±0.02 |
| LPC | lysophosphatidylcholine | 0.08%±0.01 | 0.06%±0.01 | 0.13%±0.04 | 0.07%±0.02 | 0.09%±0.05 | 0.07%±0.02 |
| PG | phosphatidylglycerol | 0.06%±0.01 | 0.03%±0 | 0.05%±0.01 | 0.08%±0.01 | 0.06%±0.01 | 0.06%±0.01 |

[1]SM refers to a sphingomyelin with a choline group

**Table S8** The molar proportions of each fatty acid methyl ester (FAME) found in each treatment group. Values are grouped means (averaged across the six replicates of each treatment group) +/- two standard errors.

| FAME Species | ND Before | ND Warm 15-d | ND Cold 15-d | D Before | D Warm 15-d | D Cold 15-d |
|---|---|---|---|---|---|---|
| C18.1n9c | 27.83%±0.29 | 28.11%±0.41 | 27.4%±0.43 | 27.77%±0.71 | 27.72%±0.47 | 27.39%±0.36 |
| C16.0 | 25.35%±0.42 | 24.58%±0.21 | 24.73%±0.29 | 25.07%±0.17 | 24.66%±0.28 | 24.87%±0.23 |
| C16.1n7 | 21.04%±1.12 | 21.08%±1.07 | 18.77%±0.93 | 20.36%±2.53 | 19.61%±2.01 | 18.46%±1.4 |
| C18.2n6c | 14.81%±0.55 | 14.86%±0.65 | 16.05%±0.61 | 14.41%±0.91 | 14.78%±0.67 | 15.32%±0.17 |
| C18.0 | 3.86%±0.39 | 4.16%±0.33 | 4.49%±0.25 | 4.07%±0.72 | 4.39%±0.61 | 4.54%±0.45 |
| C14.0 | 1.82%±0.15 | 1.79%±0.12 | 2.13%±0.16 | 2.29%±0.46 | 2.39%±0.37 | 2.51%±0.35 |
| C18.3n3 | 1.82%±0.1 | 1.77%±0.14 | 2.02%±0.07 | 1.85%±0.15 | 1.91%±0.11 | 2.02%±0.03 |
| C20.4n6 | 1.06%±0.23 | 1.25%±0.19 | 1.46%±0.16 | 1.1%±0.29 | 1.27%±0.27 | 1.37%±0.17 |
| C14.1n5 | 0.68%±0.09 | 0.65%±0.05 | 0.76%±0.08 | 0.95%±0.15 | 0.95%±0.13 | 1.02%±0.12 |
| C12.0 | 0.63%±0.06 | 0.59%±0.03 | 0.72%±0.06 | 0.78%±0.18 | 0.8%±0.13 | 0.86%±0.15 |
| C20.0 | 0.35%±0.08 | 0.37%±0.06 | 0.47%±0.05 | 0.43%±0.13 | 0.48%±0.13 | 0.52%±0.09 |
| C20.5n3 | 0.34%±0.07 | 0.37%±0.05 | 0.47%±0.05 | 0.39%±0.1 | 0.44%±0.08 | 0.49%±0.05 |
| C18.3n6 | 0.14%±0.03 | 0.14%±0.02 | 0.19%±0.02 | 0.2%±0.03 | 0.22%±0.02 | 0.24%±0.01 |
| C17.0 | 0.1%±0.01 | 0.1%±0.01 | 0.12%±0.01 | 0.11%±0.02 | 0.12%±0.02 | 0.12%±0.02 |
| C15.0 | 0.09%±0.01 | 0.08%±0.01 | 0.1%±0.01 | 0.11%±0.02 | 0.11%±0.02 | 0.12%±0.01 |
| C22.0 | 0.04%±0.01 | 0.05%±0.01 | 0.07%±0.01 | 0.06%±0.03 | 0.08%±0.03 | 0.09%±0.02 |
| C20.3n6 | 0.03%±0.01 | 0.02%±0.01 | 0.03%±0.01 | 0.04%±0.01 | 0.04%±0.01 | 0.05%±0.01 |
| C13.0 | 0.02%±0 | 0.02%±0 | 0.02%±0 | 0.02%±0 | 0.02%±0 | 0.02%±0 |

**Table S9** Results of non-parametric Kruskal-Wallis test of whether lipid summary variables differed significantly across any of the six total treatment groups. Abbreviations are as follows: FA = fatty acid; PE = phosphoethanolamine; PC = phosphatidylcholine

| Variable | Kruskal-Wallis chi-squared | df | *p*-value |
|---|---|---|---|
| PE:PC | 14.664 | 5 | 0.012 |
| FA Unsaturation Index | 20.327 | 5 | 0.001 |
| Proportion Poly-unsaturated FA | 13.339 | 5 | 0.020 |
| Average FA Length | 14.598 | 5 | 0.012 |

**Table S10** Results of non-parametric post-hoc testing (pairwise Wilcox tests) comparing lipid summary variables across all pairwise comparisons of diapause and acclimation treatments. Treatment abbreviations are as follows: ND = non-diapause; D = diapause; Before = before acclimation; Warm = warm acclimation; Cold = cold acclimation; Deacc = de-acclimation. Significant *p*-values are bolded.

| Contrast | PE:PC *p*-value | FA Unsaturation Index *p*-value | Proportion Poly-unsaturated FA *p*-value | Average FA Length *p*-value |
|---|---|---|---|---|
| D Cold 15-d – D Before | 0.937 | **0.013** | 0.248 | 0.162 |
| D Warm 15-d – D Before | 0.283 | 0.139 | 0.736 | 0.464 |
| D Warm 15-d – D Cold 15-d | 0.283 | 0.198 | 0.400 | 0.631 |
| ND Before – D Before | 0.877 | 0.631 | 1.000 | 0.631 |
| ND Before – D Cold 15-d | 0.877 | 0.155 | 0.130 | 0.400 |
| ND Before – D Warm 15-d | 0.537 | 0.455 | 0.727 | 0.631 |
| ND Cold 15-d – D Before | 0.400 | **0.011** | 0.139 | **0.043** |
| ND Cold 15-d – D Cold 15-d | 0.337 | **0.013** | 0.139 | **0.043** |
| ND Cold 15-d – D Warm 15-d | 0.736 | **0.011** | 0.139 | 0.057 |
| ND Cold 15-d – ND Before | 0.464 | **0.011** | **0.032** | **0.043** |
| ND Warm 15-d – D Before | **0.016** | 0.155 | 1.000 | 0.162 |
| ND Warm 15-d – D Cold 15-d | **0.016** | 0.937 | 0.139 | 0.699 |
| ND Warm 15-d – D Warm 15-d | 0.195 | 0.455 | 1.000 | 0.631 |
| ND Warm 15-d – ND Before | 0.076 | 0.328 | 0.736 | 0.400 |
| ND Warm 15-d – ND Cold 15-d | 0.097 | **0.038** | 0.114 | 0.283 |
